# Specific interaction between Group B *Streptococcus* CC17 hypervirulent clone and phagocytes

**DOI:** 10.1101/2022.12.02.518834

**Authors:** Anne-Sophie Bourrel, Amandine Picart, Jose-Carlos Fernandez, Constantin Hays, Virginie Mignon, Bruno Saubaméa, Claire Poyart, Agnès Fouet, Asmaa Tazi, Julie Guignot

## Abstract

*Streptococcus agalactiae* also named Group B *Streptococcus* (GBS) is the most significant pathogen causing invasive infections, such as bacteremia and meningitis, in neonates. Worldwide epidemiological studies have shown that a particular clonal complex (CC) of capsular serotype III, the CC17, is strongly associated with meningitis in neonates and is therefore designated as the hypervirulent clone. Macrophages are a permissive niche for intracellular bacteria of all GBS clones. In this study we deciphered the specific interaction of GBS CC17 strains with macrophages. Our study revealed that CC17 strains are phagocytosed at a higher rate than GBS non-CC17 strains by human monocytes and macrophages both in cellular models and primary cells. CC17 enhanced phagocytosis is due to an initial enhanced-attachment step to macrophages mediated by the CC17 specific surface protein HvgA and the PI-2b pilus (Spb1). We showed that two different inhibitors of scavenger receptors (fucoidan and poly(I)) specifically inhibited CC17 adhesion and phagocytosis while not affecting those of non-CC17 strains. Once phagocytosed, both CC17 and non-CC17 strains remained in a LAMP-1 positive vacuole that ultimately fuses with lysosomes where they can survive at similar rates. Finally, both strains displayed a basal egress which occurs independently from actin and microtubule networks. Our findings provide new insights into the interplay between the hypervirulent GBS CC17 and major players of the host’s innate immune response. This enhanced adhesion leading to higher phagocytosis could reflect a peculiar capacity of the CC17 lineage to subvert the host immune defenses, establish a niche for persistence or disseminate.

## INTRODUCTION

The innate immune system plays a crucial role during host’s interaction with pathogens, particularly in neonates who exhibit a naive adaptive immune system and are more vulnerable to invasive infections (1). Innate immunity largely relies on professional phagocytic cells, which constitute the first line of defense against pathogens. The large repertoire of phagocytic surface receptors that mediate the binding and internalisation of the pathogen is divided in opsonic and non-opsonic receptors. Opsonic receptors include the Fc receptors and complement receptors, which recognize antibody- or complement-opsonized pathogens, but also some non-opsonized pathogens (2, 3). Non-opsonic receptors are constituted by a wide array of family receptors including Toll like receptors (TLRs), integrins, C-type lectins or scavenger receptors that directly interact with pathogen-associated molecular patterns (PAMPs). Receptor recognition triggers intracellular signaling cascades leading to actin polymerization-which generates membrane protrusions required for bacterial engulfment. Signal transduction during phagocytosis is specific to the cell type and to the receptor(s) involved and can impact the intracellular fate of the microorganism (4). Once internalized, the pathogen resides in a phagosome, which rapidly undergoes maturation and acquires numerous proteins such as the late endosome/lysosome-associated membrane proteins LAMP-1 and LAMP-2. Ultimately, the phagosome fuses with lysosomes to become a phagolysosome with microbicidal features including a very acidic lumen (pH ∼ 4.5), lysosomal proteases, and reactive oxygen species (ROS). This in turn leads to microbial killing and production of pro-inflammatory mediators that orchestrate the initiation of adaptive immunity.

*Streptococcus agalactiae*, also named Group B *Streptococcus* (GBS), is a capsulated chain-forming Gram-positive coccus commonly found in the gut and the lower urogenital tract of approximately 20% healthy adults (5). Although rarely pathogenic in adults, GBS is the leading cause of invasive infections in neonates (6). Two neonatal GBS-associated syndromes, referred to as “early-onset disease” (EOD) and “late-onset disease” (LOD), have been described, depending on the age at onset. EOD often manifests by respiratory distress that rapidly progresses into bacteremia and less frequently meningitis. In contrast, LOD manifests as a bacteremia highly associated with meningitis (20-50% cases) (7, 8). Based on the capsular polysaccharide composition, GBS has been classified into 10 distinct serotypes (Ia, Ib, and II–IX). Serotype III is the most frequently identified serotype in young infants (61.5%) (9). More specifically, within serotype III, multilocus sequence typing (MLST) has identified a hypervirulent lineage termed clonal complex 17 (CC17) which accounts for a substantial proportion (25%) of EOD cases and the vast majority (70%) of LOD cases worldwide, whereas it is isolated in about only 13% of non-pregnancy-related adult infections and colonization (7, 10, 11). GBS CC17 is a highly successful invasive clone in the neonate strongly associated with neonatal meningitis (∼70%) in both EOD and LOD (12–14). CC17 strains harbor a number of specific genes including the genes encoding two surface proteins named Srr2 and HvgA that are involved in the crossing of the intestinal and blood brain barriers (8, 15–17). Although not totally exclusive to the CC17 lineage, the pilus island PI-2b encoding the Spb1 adhesin is found in the vast majority of the CC17 strains, while rarely present in other human lineages (15, 18).

GBS has developed several mechanisms to resist immune detection, phagocytosis and microbial degradation (19). The capsular polysaccharide (CPS) is one of the major virulence factors of GBS allowing evasion from the host immune system, notably by preventing host complement deposition on the bacterial surface (20, 21). Although capsule limits phagocytosis of complement-opsonized GBS, opsonized-GBS can be phagocytosed by Fc and complement receptors respectively (19, 22). However, there are large gaps in our understanding of the phagocytosis of non-opsonized GBS. The receptor(s) and signaling pathway(s) involved in non-opsonized GBS phagocytosis are unknown. Interestingly, non-opsonized GBS uptake leads to a better survival within phagocytes (19, 23–25). The persistence of GBS in macrophages relies on numerous bacterial factors (19, 26). The expression of some virulence factors associated to the CC17 lineage such as Spb1 have been described to increase phagocytosis (27–29). However, the capacity of the CC17 lineage to exhibit enhanced phagocytosis and survival in macrophages compared to those of other GBS lineages is still disputed (25, 30–32).

In the present study, we used a collection of GBS isolates representing a range of capsular types and CCs and showed that CC17 strains are more phagocytosed than non-CC17 isolates. Our data support that the enhanced phagocytosis of CC17 strains is solely due to an increased initial adhesion to phagocytes. Once internalized, GBS CC17 and non-CC17 do not differentially affect the phagolysosome maturation nor their ability to survive within phagocytes and can be released in the extracellular compartement.

## RESULTS

### GBS CC17 strains are more phagocytosed than non-CC17 strains by macrophages and monocytes

The literature shows conflicting results on GBS lineages capacity to be phagocytosed (25, 30–32). Therefore, we performed a phagocytosis assay using THP-1 differentiated macrophages to compare the level of phagocytosis of a relevant collection of GBS isolates responsible for human invasive infections including 27 CC17 strains and 40 non–CC17 strains (Table S2, Fig. 1A). The CC17 strains were more phagocytosed than the non-CC17 strains (2.3-fold increase). To further characterize the lineage differences and the phagocytosis process, we next used two well-characterized and fully sequenced GBS isolates, one CC17 strain (BM110) and one CC23 strain (NEM316). They displayed a difference in phagocytosis level comparable to that of CC17 versus non-CC17 strains in THP-1 macrophages (2.5-fold increase) (Fig. S1). Using these two strains, we confirmed the higher phagocytosis of GBS CC17 in primary human monocytes derived macrophages (hMDM) (Fig. 1B), in non-differentiated THP-1 monocytes (Fig. 1C) and in primary human monocytes (Fig. 1D).

**Figure 1:**
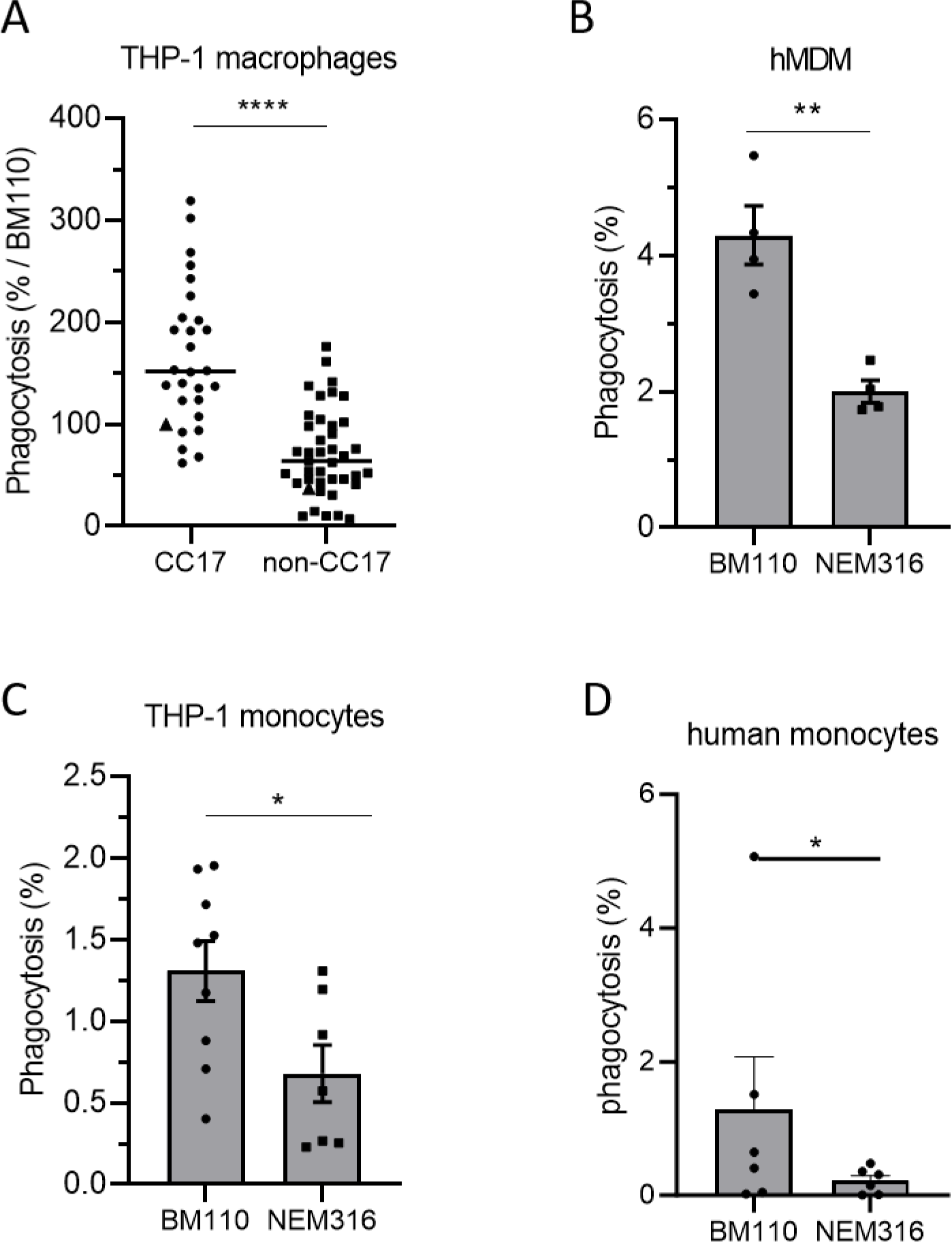
CC17 GBS strains are more phagocytosed by monocytes and macrophages than non-CC17 GBS strains. (A, B, C, D) Phagocytosis level of GBS strains was assessed by CFU count after infection at MOI 10 followed by antibiotic treatment to kill extracellular bacteria using (A) THP-1 macrophages (B) hMDM primary cells, (C) THP-1 monocytes or (D) human primary monocytes. (A) Phagocytes were infected with CC17 and non-CC17 GBS clinical isolates from invasive infections, each dot representing a clinical strain. Triangles correspond to BM110 strain (CC17) and NEM316 (non-CC17). Results are expressed as the percentage of phagocytosis relative to BM110 strain phagocytosis, with horizontal lines indicating median value. (B, C, D) Phagocytes were infected with GBS strains BM110 and NEM316. Results are expressed as the percentage of phagocytosed bacteria normalized to (B, C) the initial inoculum or (D) to BM110 strain phagocytosis. Statistical analysis: data shown are (A) median or (B, C, D) mean ± SEM of at least four independent experiments. (A, B, C) t test or (D) Mann Whitney test, were performed with *, *p* < 0.05; **, *p* < 0.01; ****, *p* < 0.0001.

Since the capsule is a major bacterial factor involved in GBS phagocytosis, we analyzed its contribution to the enhanced phagocytosis of GBS CC17 strains. First, we stratified the results of the phagocytosis assay performed with the clinical GBS strains presented in Fig. 1A according to their capsular type. CC17 strains being almost exclusively of capsular type III and more rarely of type IV, we distinguished within CC17 and non-CC17 isolates those of capsular type III, IV and all the other types (Fig. 2A). The CC17 strains higher phagocytosis was independent from the capsular type. To further test the influence of the capsular type, we analyzed the phagocytosis level of BM110 and NEM316 mutants, inactivated for capsule expression. As expected, an increased phagocytosis of the capsule deficient strains was observed with both mutant strains (Fig. 2B). Furthermore, the higher phagocytosis of the CC17 isolate compared to the non-CC17 isolate could also be observed with the corresponding non-capsulated mutants (Fig. 2B).

**Figure 2:**
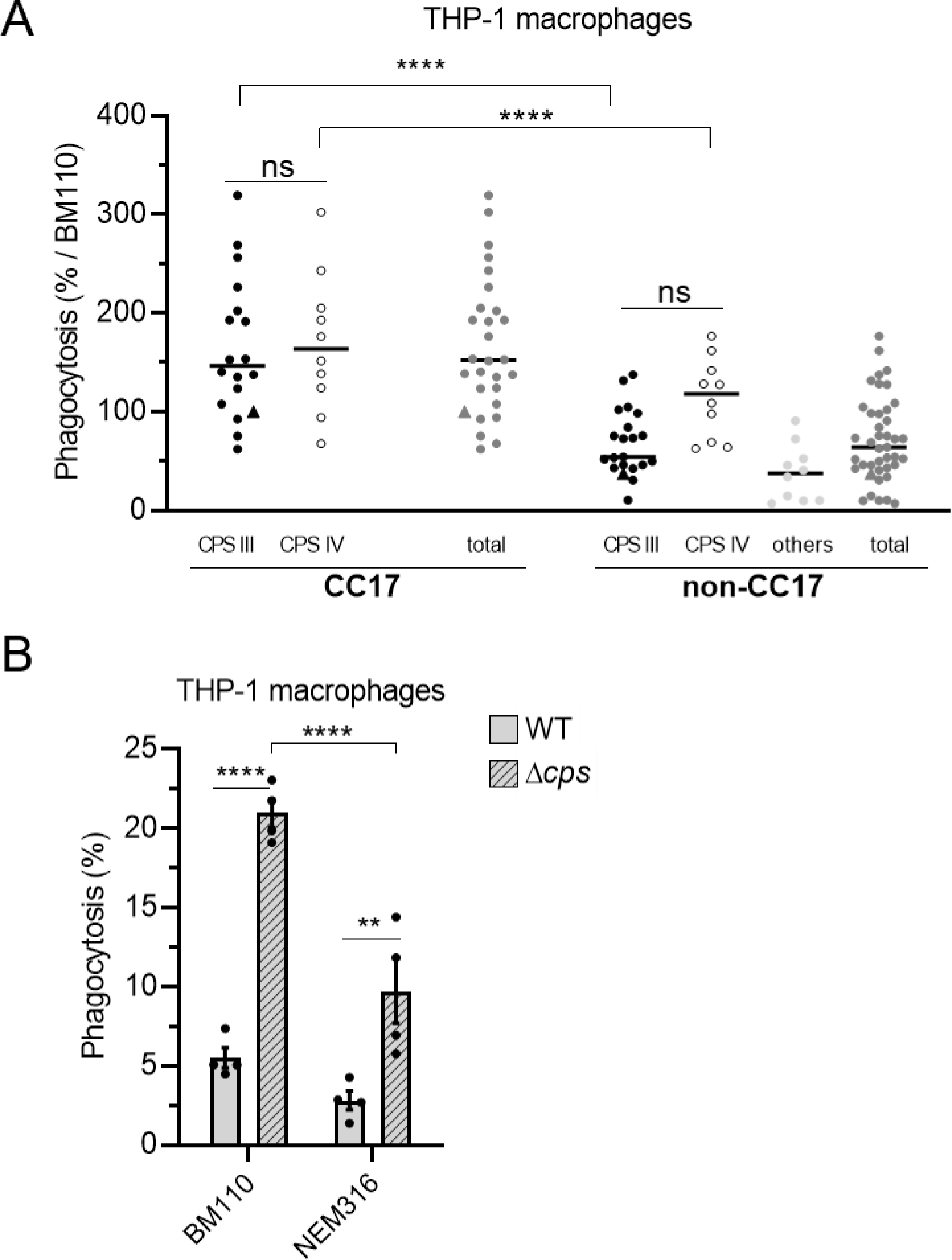
GBS capsule is not involved in the hyper phagocytosis of CC17 strains. (A, B) Phagocytosis level of GBS strains was assessed by CFU count after infection at MOI 10, followed by antibiotic treatment to kill extracellular bacteria. (A) THP-1 macrophages were infected with CC17 and non-CC17 GBS clinical isolates from invasive infections, each dot representing a clinical strain. Triangles correspond to BM110 strain (CC17) and NEM316 (non-CC17). Results are shown according to capsular type (CPS III, CPS IV, or other CPSs) and are expressed as the percentage of phagocytosis relative to BM110 strain phagocytosis, with horizontal lines indicating median value. (B) THP-1 macrophages were infected with GBS strains BM1110 and NEM316 and their respective mutant strains impaired for capsule expression (Δ*cps*). Results are expressed as the percentage of phagocytosed bacteria normalized to the initial inoculum. Statistical analysis: data shown are (A) median or (B) mean ± SEM of at least four independent experiments. (A, B) Two-way ANOVA, was performed with ns, non-significant; **, *p* < 0.01; ****, *p* < 0.0001.

Altogether, these results show that CC17 strains are more phagocytosed than non-CC17 strains in a capsule independent manner.

### HvgA and Spb1 promote phagocytosis of CC17 strains

This observation prompted us to search for CC17 specific virulence factors that may account for this enhanced phagocytosis. To this end, we tested the phagocytosis level of BM110 inactivated in genes encoding CC17-specific (Srr2, HvgA) or -highly associated (Spb1) surface proteins (Fig. 3A) (15). Both *ΔhvgA and Δspb1* mutants exhibited reduced phagocytosis in THP-1 macrophages (65% and 54% reduction, respectively) compared to the wild-type (WT) strain. In contrast, there was no difference between the phagocytosis of the Δ*ssr2* and the WT strain. The role of Spb1 in GBS phagocytosis has already been described (27, 28, 33), but not that of HvgA. To confirm the direct involvement of the HvgA surface protein in phagocytosis, we used a close relative non-pathogenic *Lactococcus lactis*, in which an expression vector encoding HvgA and enabling its presentation at the bacterial surface was introduced (16). The heterologous expression of HvgA in *L. lactis* led to a 3-fold increase in phagocytosis, as compared to *L. lactis* carrying an empty vector (Fig. 3B). The involvement of HvgA in CC17 phagocytosis was also observed using hMDM (Fig. 3C).

**Figure 3:**
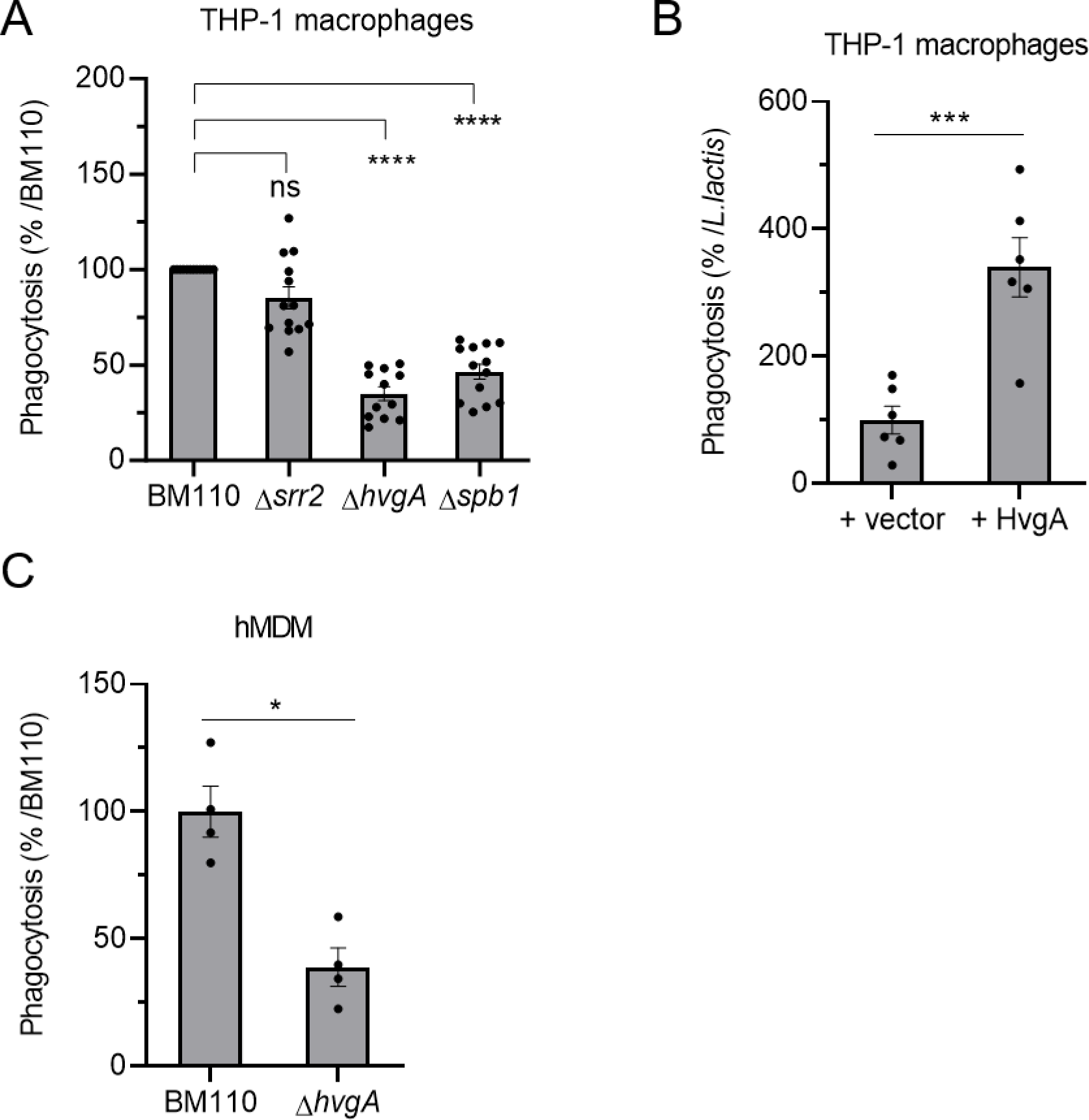
The HvgA surface protein and Spb1 pilus protein are involved in CC17 hyper-phagocytosis. (A, B, C) Phagocytosis level of GBS strains was assessed by CFU count after infection at MOI 10 followed by antibiotic treatment to kill extracellular bacteria using (A, B) THP-1 macrophages (C) hMDM primary cells. (A, C) Macrophages were infected with BM110 and its derivative mutant strains (Δ*srr2*, Δ*hvgA* or Δ*spb1*), or (B) with *Lactococcus lactis* strain carrying an empty vector (+vector) or a vector expressing the HvgA protein (+HvgA). (A, C) Results are expressed as the relative levels to BM110 strain, or (B) to the control *Lactococcus lactis* strain +vector. Statistical analysis: data shown are mean ± SEM of at least four independent experiments. (A) Kruskall Wallis test with Dunn’s multiple comparisons or (B, C) t test, were performed with ns, non-significant; *, *p* < 0.05; ***, *p* < 0.001; ****, *p* < 0.0001.

### The signal transduction leading to phagocytosis is similar between CC17 and non-CC17 strains

Cell membrane protrusion formation leading to bacterial engulfment results from cytoskeleton rearrangements, which are mediated by specific signaling cascades induced by bacterial binding to cell surface receptors (34). Numerous signaling molecules regulate the cytoskeleton, including the small Rho GTPases Rho, Rac and CDC42 known to allow the formation of actin stress fibers, lamellipodial protrusions, and filopodia, respectively (35). To better characterize the molecular pathways involved in GBS engulfment, we performed phagocytosis assays using a wide array of inhibitors targeting different signaling molecules (Table S3, Fig. 4A). BM110 phagocytosis was inhibited by cytochalasin D, ZCL278, staurosporine, LY29002 and PP2 indicating that CC17 phagocytosis required actin, the small GTPase CDC42, as well as the different kinases, PKC, Pi3K and Src, respectively. In contrast, microtubules, the small GTPase Rac and the kinases Syk and Rock were not involved in CC17 phagocytosis, as indicated by the absence of effect of nocodazole, NSC23766, BAY61-3606 and Y27632 respectively (Fig. 4A). Interestingly, we found that NEM316, BM110Δ*hvgA* or BM110Δ*spb1* phagocytosis involved the same signaling proteins than BM110 (Fig. 4B, 4C and 4D respectively).

**Figure 4:**
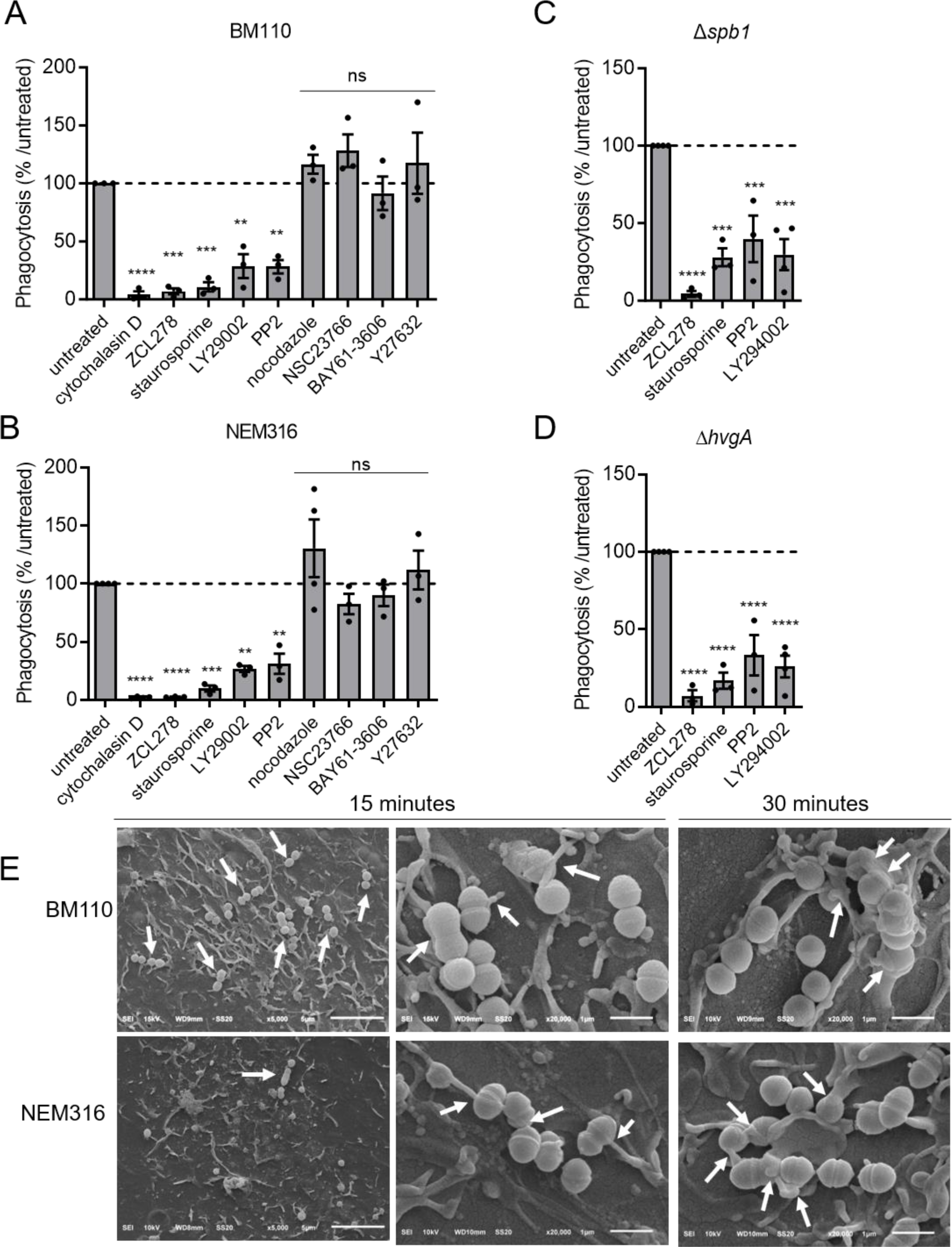
Signal transduction leading to phagocytosis is similar for CC17 and non-CC17 strains. (A, B, C, D) THP-1 macrophages were pre-treated for one-hour prior infection with cytochalasin D, ZCL178, staurosporin, LY29002, PP2, nocodazole, NSC23766, BAY61-3606 or Y27632 to inhibit actin polymerization, CDC42, protein kinase C, PI3kinase, Src kinase, microtubule polymerization, Rac, Syc and Rock, respectively. (A) Phagocytosis level of BM110 (B) NEM316 (C) Δ*spb1* or (D) Δ*hvgA* GBS strains were assessed by CFU count after infection at MOI 10 followed by antibiotic treatment to kill extracellular bacteria. Results are expressed as the relative levels to untreated condition. (E) Scanning electron microscopy (SEM) micrographs of THP-1 macrophages showing (left panel, scale bar 5 µm) bacterial adherence to cell surface and (middle panel and right panels, scale bar 1 µm) bacterial capture by filopodia after 15 min or lamellipodia after30 min of infection. Statistical analysis: data shown are mean ± SEM of at least four independent experiments. One-Way ANOVA with Dunnett multiple comparisons, were performed with ns, non-significant; **, *p* < 0.01; ***, *p* < 0.001, ****, *p* < 0.0001.

To gain further insight into these findings, we characterized the early contact between GBS and macrophages, which occurs prior to phagocytosis by Scanning Electron Microscopy (SEM) (Fig. 4E). SEM revealed that numerous BM110 chains were adherent to macrophages in contrast to only a few NEM316 chains (Fig. 4E, left panel). In addition, the uptake of both BM110 and NEM316 was accompanied by the formation of filopodia-like cellular protrusions that were observed at 15 min (Fig.4E, middle panel) and lamellipodia-like structure at 30 min (Fig. 4E, right panel).

Taken together, our results show that CC17 and non-CC17 are both captured by filopodia and that bacterial uptake involves similar signaling proteins. Our results also suggest that enhanced phagocytosis of CC17 strains may be due, at least in part, to an increased adhesion.

### CC17 enhanced phagocytosis is due to initial higher adhesion

To test a potential adhesion difference between CC17 and non-CC17 strains, we analyzed the adhesion of GBS strains to THP-1 macrophage. To prevent bacterial engulfment, adhesion was assessed either at 4°C or at 37°C in the presence of cytochalasin D, an inhibitor of actin polymerization (Fig. S2A). Data showed that bacterial adhesion to phagocytes was similar for all strains independently of the method used to block phagocytosis. In both cases, adhesion of the BM110 strain was higher than that of NEM316. We next assessed adhesion at 4°C and observed a higher adhesion of CC17 clinical strains than of non-CC17 strains (1.9-fold increase, Fig. 5A, Table S2) to THP-1 macrophages. The enhanced adhesion was also observed with strain BM110 compared to NEM316 on THP-1 macrophages (Fig. S2A), THP-1 monocytes (Fig. 5B) and hMDM (Fig. 5C). As observed for phagocytosis, the increased adhesion of CC17 isolates was not related to the capsular serotype (Fig. S2B) and was still observed in the absence of capsule expression (Fig. 5D). Both *ΔhvgA and Δspb1* mutants exhibited reduced adhesion on THP-1 macrophages (23% and 48% reduction, respectively) compared to the WT strain (Fig. 5E). The heterologous expression of HvgA in *L. lactis,* led to a 2-fold increase in adhesion to THP-1 macrophages, relative to *L. lactis* carrying an empty vector (Fig. 5F). The involvement of HvgA in CC17 adhesion was also observed using hMDM (Fig. S2C). These results indicate that CC17 strains have a higher level of adhesion compared to non-CC17 strains. Furthermore, the increased adhesion of CC17 strains is mediated by both Spb1 and HvgA.

**Figure 5:**
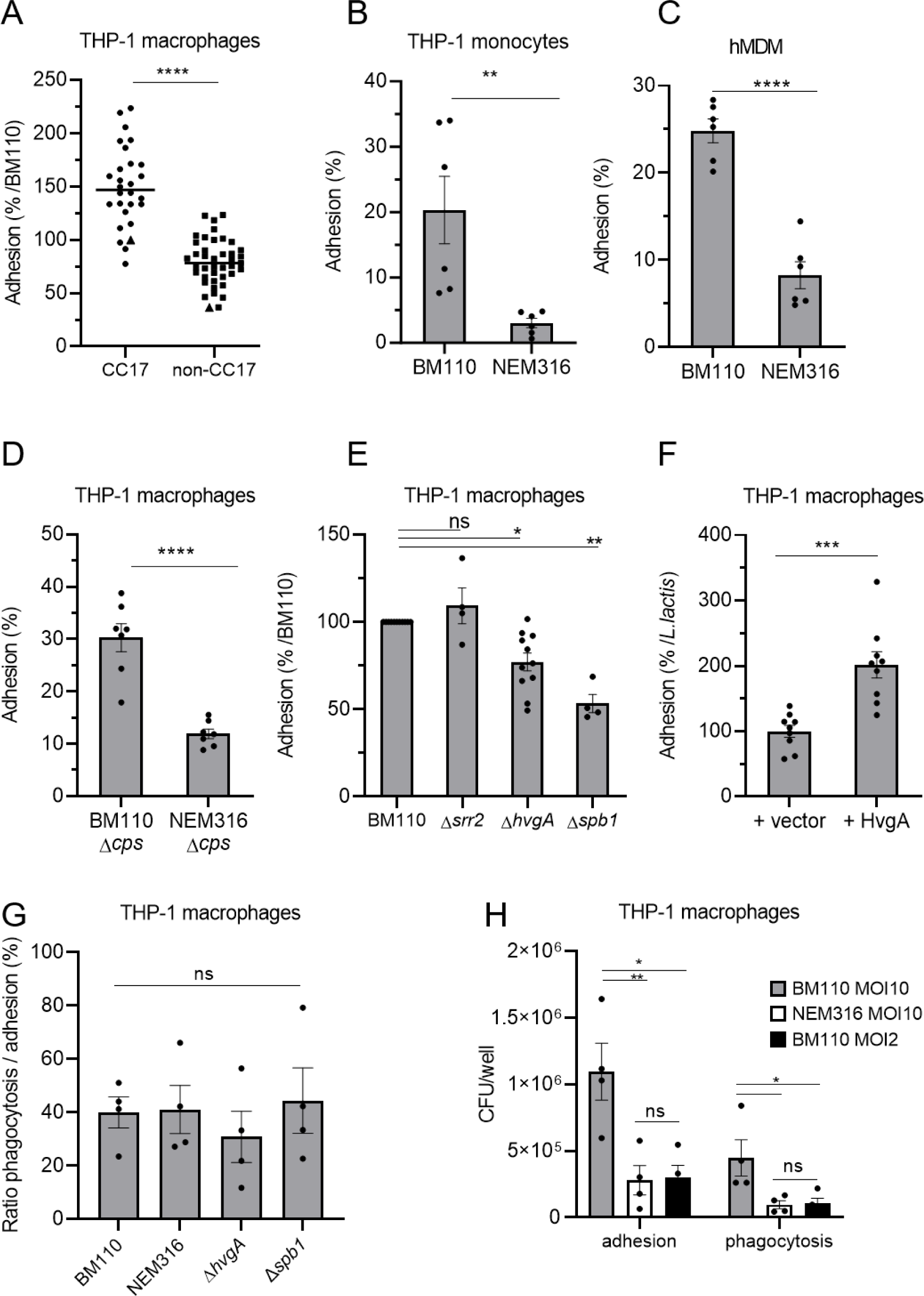
CC17 GBS strains are more adherent to phagocytes than non-CC17 strains. (A, B, C, D, E, F) Adhesion level of GBS strains was assessed by CFU count after infection at MOI 10 at 4°C to avoid bacterial engulfment using (A, D, E, F) THP-1 macrophages (B) THP-1 monocytes (C) hMDM primary cells. (A) Phagocytes were infected with CC17 and non-CC17 GBS clinical isolates from invasive infections, each dot representing a clinical strain. Triangles correspond to BM110 strain (CC17) and NEM316 (non-CC17). Results are expressed as the percentage of adhesion relative to BM110 strain adhesion, with horizontal lines indicating median value. (B, C, D, E, G, H) Phagocytes were infected with GBS strains or (F) with *Lactococcus lactis* strain carrying an empty vector (+vector) or a vector expressing HvgA protein (+HvgA). Results were normalized to (B, C, D) the initial inoculum (E) BM110 strain or (F) *L. lactis* control strain (+vector). (G,H) (G) Adhesion and phagocytosis were assessed by CFU count after infection at MOI 10. Results were expressed as the ratio between CFU of phagocytosed bacteria and adherent bacteria. (H) Adhesion and phagocytosis were assessed by CFU count after infection at MOI 10 or 2. Results were normalized to the initial inoculum. Statistical analysis: data shown are mean ± SEM of at least four independent experiments. (A, B, C, D, F) t Test (E) Kruskall Wallis test with Dunn’s multiple comparisons or (G, H) One-Way ANOVA with Tukey’s multiple comparisons test, were performed with ns, non-significant; *, *p* < 0.05; **, *p* < 0.01; ***, *p* < 0.001; ****, *p* < 0.0001.

To determine whether CC17 higher phagocytosis was solely due to the increased adhesion or to a more complex phenomenon involving both increased adhesion and uptake, we compared the ratio of phagocytosed bacteria to the initial number of adherent bacteria. BM110 and its derivative mutant strains and NEM316 strains showed similar ratios of phagocytosed bacteria (Fig. 5G). Next, we infected THP-1 macrophages with the strain BM110 at a lower multiplicity of infection than NEM316 (MOI 2 and MOI 10, respectively) to obtain similar numbers of adherent bacteria (Fig. 5H). In that case, similar numbers of phagocytosed bacteria were obtained with both strains.

Taken together, our results show that the enhanced phagocytosis of BM110 strain is solely due to its initial increased adhesion to macrophages.

### CC17 uptake by macrophages is inhibited by scavenger receptor inhibitors

Phagocytes express multiple receptors involved in bacterial adhesion and uptake. To identify the precise nature of the macrophage receptor(s) involved in GBS recognition and uptake, GBS CC17 phagocytosis was evaluated in cells pre-treated with various blocking molecules targeting putative receptor(s) (Table S3, Fig. 6A). Whereas the inhibitors targeting FCyR, complement receptors, integrins and lectin receptors did not affect GBS CC17 phagocytosis, fucoidan and poly(I), two different scavenger receptor inhibitors, significantly reduced it by 60% and 32%, respectively. Importantly, the inactive analog of poly(I), poly(C), used as a negative control, had no effect on CC17 phagocytosis (Fig. 6A). Interestingly, these phenotypes were specific to BM110 since no reduction of NEM316 phagocytosis was observed using these two scavenger receptor inhibitors (Fig. 6B). Importantly, fucoidan was also found to reduce BM110 adhesion to THP-1 macrophages whereas it did not reduce NEM316 adhesion (Fig. 6C). These results suggest that some scavenger receptors contribute to a significant proportion of the GBS CC17 adhesion that lead to its uptake into macrophages.

**Figure 6:**
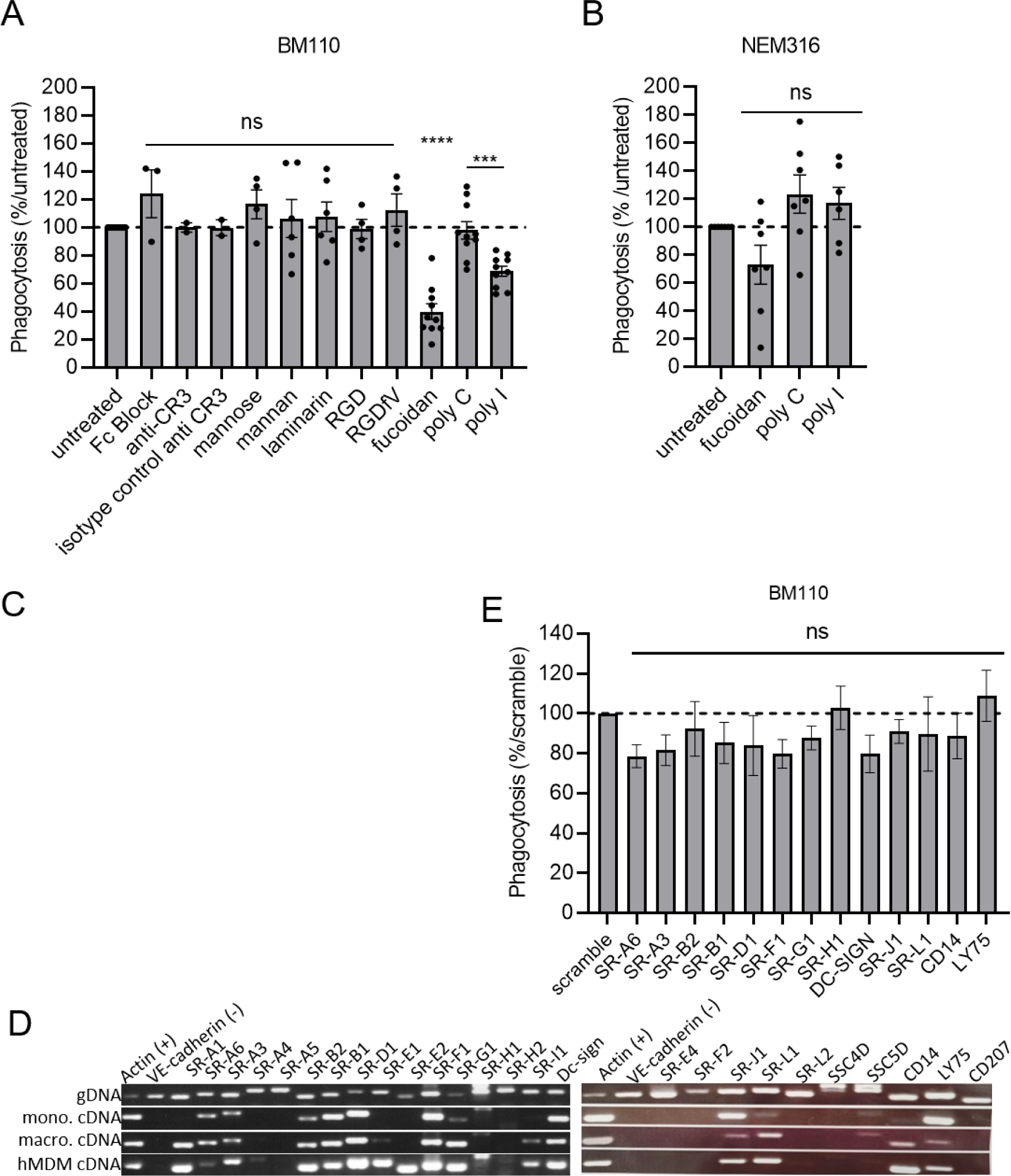
Scavenger receptor inhibitors lead to a decreased phagocytosis level of CC17 GBS in macrophages. (A, B, E) Phagocytosis level in THP-1 macrophages was assessed by CFU count after infection at MOI 10 with (A, E) BM110 or (B) NEM 316 GBS strains followed by 1h antibiotic treatment to kill extracellular bacteria. (A) Macrophages were pre-treated 1h before infection to block FCγ receptors (Fc Block™), complement receptor 3 (CR3 blocking antibody), lectin receptors (mannose, mannan, laminarin), integrins receptors (RGD and RGDfV mimetic peptides), or (A, B) scavenger receptors inhibitors (Fucoidan and poly(I) or inactive analog poly(C). (A, B) Results are expressed as the percentage of phagocytosis relative to the untreated control condition except for poly (I) that was relative to its inactive analogue poly(C). (C) Adhesion level in untreated or fucoidan treated-THP-1 macrophages was assessed by CFU count after infection with BM110 and NEM316 at 4°C to avoid bacterial engulfment. Results are expressed as the percentage of phagocytosis relative to the untreated control condition (D) Expression of scavenger receptors genes was assessed by RT-PCR using THP-1 macrophages (macro), THP-1 monocyte (mono) and hMDM (hMDM) cDNA. Actin and VE-cadherin gene expression was used as positive and negative controls of expression, respectively. Genomic DNA (gDNA) was used to validate primers efficiency. (E) Involvement of scavenger receptors on BM110 phagocytosis was assessed by silencing scavenger receptors expression by si-RNA. Results are expressed as the percentage of phagocytosis relative to the scramble control condition. Statistical analysis: data shown are mean ± SEM of at least four independent experiments. (A, B, E) Kruskall Wallis test with Dunn’s multiple comparisons or (C) t test were performed, with ns, non-significant; **, *p* < 0.01; ***, *p* < 0.001; ****, *p* < 0.0001.

Scavenger receptors constitute a large family of membrane-associated pattern recognition receptors (PRRs) that act as receptors mediating direct non-opsonic adhesion and uptake of microbes and/or their products (36). To determine which scavenger receptor(s) contribute(s) to the adhesion/uptake of GBS CC17 in THP-1 macrophages, we characterized by RT-PCR their expression profile in THP-1 macrophages. Given that the enhanced phagocytosis of GBS CC17 is conserved in hMDM and monocytes (Fig. 1B, C, D), we extended the analysis of scavenger receptors expression profile to these cell types (Fig. 6D). Using the recent classification of scavenger receptors (37), we identified that, out of the 25 different scavenger receptors tested, 13 were expressed by these three cellular models (Fig. 6D). Next, we silenced the expression of each of these 13 individual scavenger receptors by siRNA to address their involvement in GBS CC17 uptake. Efficient knockdown levels were confirmed on two individual genes expressed by THP-1 macrophages for which the knockdown efficiency was 78 and 98 % (Fig. S3A). As an additional control, the silencing of the complement receptor impaired the phagocytosis of complemented-red blood cells (96% decreased) (Fig. S3B), demonstrating that siRNA in THP-1 macrophages was efficient. However, when individual scavenger receptors were silenced, no significant reduction in BM110 uptake was observed (Fig. 6E). Last, in agreement with the phagocytosis assays using phagocytosis receptor inhibitors (Fig. 6A), BM110 phagocytosis was not decreased in THP-1 macrophages silenced for FCyR, complement receptors, α5 or β3 integrin expression, Plaur or CD44 (Fig. S3C) or TLRs (Fig. S3D).

### CC17 and non-CC17 strains display similar capacity to survive within macrophages

After phagocytosis, the phagocytic vacuole containing bacteria undergoes maturation events leading to the formation of a phagolysosome required for efficient bacterial degradation. GBS is known to survive within macrophages for up to 24 hours (24, 25, 27, 31, 32). In order to gain insight into the physiological relevance of our study, we compared the intracellular fate of BM110 and NEM316 strains in THP-1 macrophages. The maturation of phagosomes containing BM110-GFP and NEM316-GFP (Table S1) was followed by fluorescence microscopy at various time-points after phagocytosis (t=0, t=2h30 and t=5h). BM110 bacteria colocalizing with LAMP-1 and with the Lysotracker (a fluorescent dye labeling acidic organelles such as phagolysosome) were quantified after immunofluorescence staining (Fig. 7A and C). No statistical differences were observed in the kinetic of phagosome maturation between BM110 and NEM316 whatever the time-point (Fig. 7B and D), indicating that the delivery of BM110 and NEM316 to late endosomes or lysosomes is similar.

**Figure 7:**
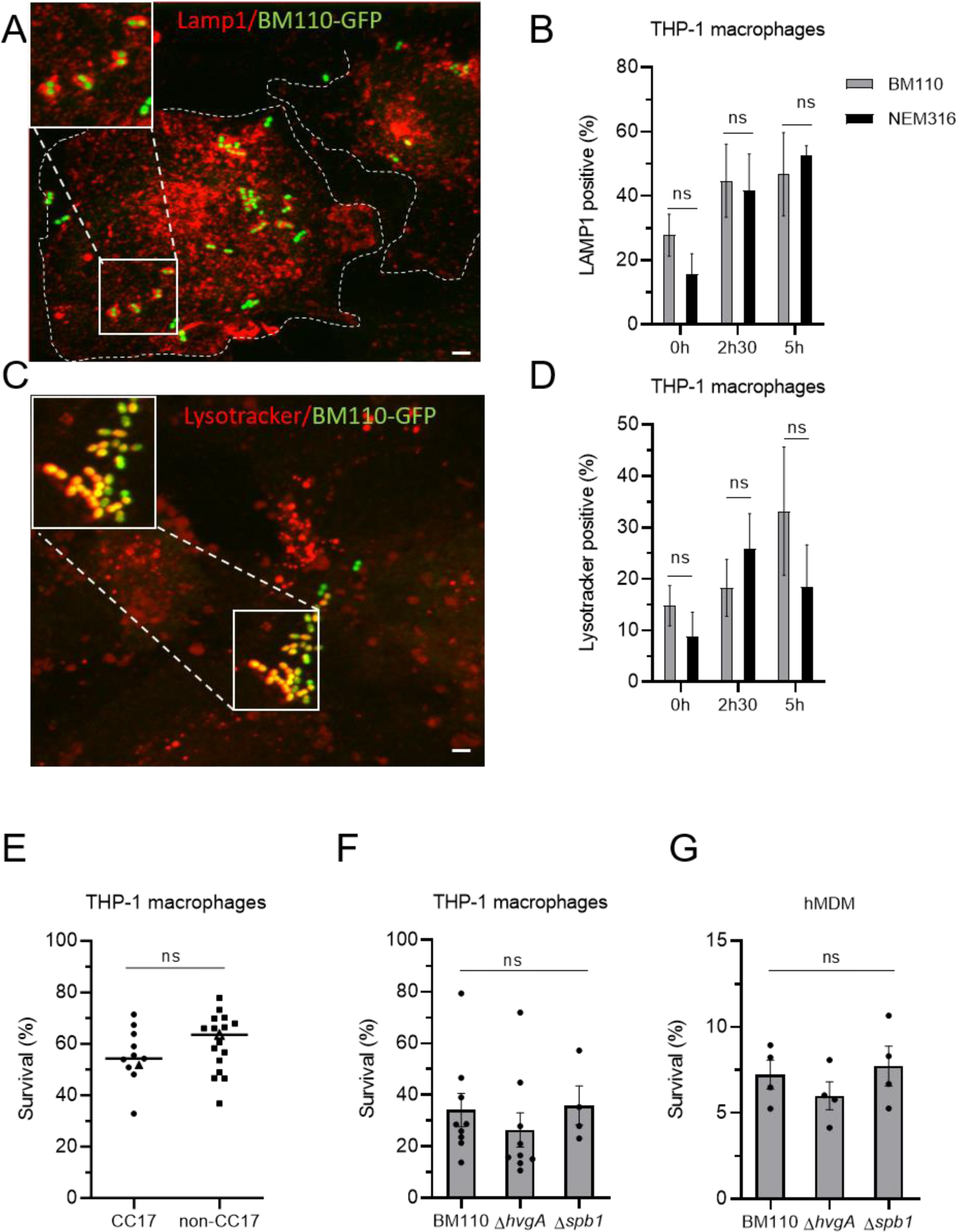
CC17 and non-CC17 GBS strains exhibit similar intracellular fate. (A, B, C, D) Intracellular fate of BM110 and NEM316 strains was followed by microscopy in THP-1 macrophages. (A, C) Representative confocal microscopy images of THP-1 macrophages infected with BM110-GFP (green) and labelled with (A) anti-LAMP1 antibody (red) or (C) Lysotracker® (red). Boxed areas correspond to magnification of insets. Dotted lines indicate cell perimeters. Scale bar: 5 µm. (B, D) Quantification of BM110-GFP and NEM316-GFP associated with (B) LAMP1 or (D) Lysotracker® was performed at 0, 2h30 or 5h post-phagocytosis. Results are expressed as the percentage of bacteria positive for (B) LAMP1 or (D) Lysotracker® staining. (E, F, G) Percentage of bacterial survival was assessed by CFU count 2h30 after phagocytosis in (E, F) THP-1 macrophage or (G) hMDM. (E) Macrophages were infected with CC17 and non-CC17 GBS clinical isolates from invasive infections, each dot representing a clinical strain. For each strain, results are expressed as the percentage of survival relative to corresponding the corresponding level of phagocytosis, with horizontal lines indicating median value. Triangles correspond to BM110 strain (CC17) and NEM316 (non-CC17) (F, G) THP-1 macrophages were infected with the BM110 strain and its derivative mutant strains (Δ*hvgA* or Δ*spb1*). Results are expressed as the percentage of viable intracellular bacteria normalized to the initial intracellular load (phagocytosis). Statistical analysis: data shown are mean ± SEM of at least four independent experiments. (B, D) Two-way ANOVA with Tuckey multiple comparison, (E) t Test or (F) Kruskall Wallis test with Dunn’s multiple comparisons or (G) One-Way ANOVA tests were performed with ns, non-significant.

We next compared the survival ability of 10 CC17 and 17 non-CC17 strains clinical isolates by quantifying intracellular viable bacteria 2h30 after phagocytosis (Fig. 7E). Survival results were normalized for each GBS strain to the number of initially phagocytosed bacteria. No significant difference was observed in the survival ability of CC17 and non-CC17 strains (Fig. 7E), whatever the capsular type (Fig. S4A). The survival rate of BM110 and NEM316 strains was also similar 24 h after phagocytosis (Fig. S4B). Consistently, HvgA and Spb1 were not involved in CC17 survival within THP-1 macrophages (Fig. 7F) or hMDM (Fig. 7G).

Taken together, these results revealed that the intracellular fate of GBS CC17 and non-CC17 after adhesion and phagocytosis is similar.

### CC17 and non-CC17 strains can escape macrophages by a passive phenomenon

We next analyzed the capacity of GBS strains to escape THP-1 macrophages. After phagocytosis and antibiotic treatment to eliminate extracellular bacteria, cells were incubated in antibiotic free medium for short periods of time (45 min and 1h30) to allow potential bacterial escape from infected cells without any bacterial multiplication within the extracellular media. GBS egress normalized to the percentage of initial phagocytosis showed that BM110 and NEM316 strains displayed similar capacity to escape from macrophages (Fig. 8A). GBS egress from THP-1 macrophages was neither inhibited by cytochalasin D nor by nocodazole (Fig. 8B), indicating that it is independent of actin cytoskeleton and microtubules, respectively.

**Fig 8:**
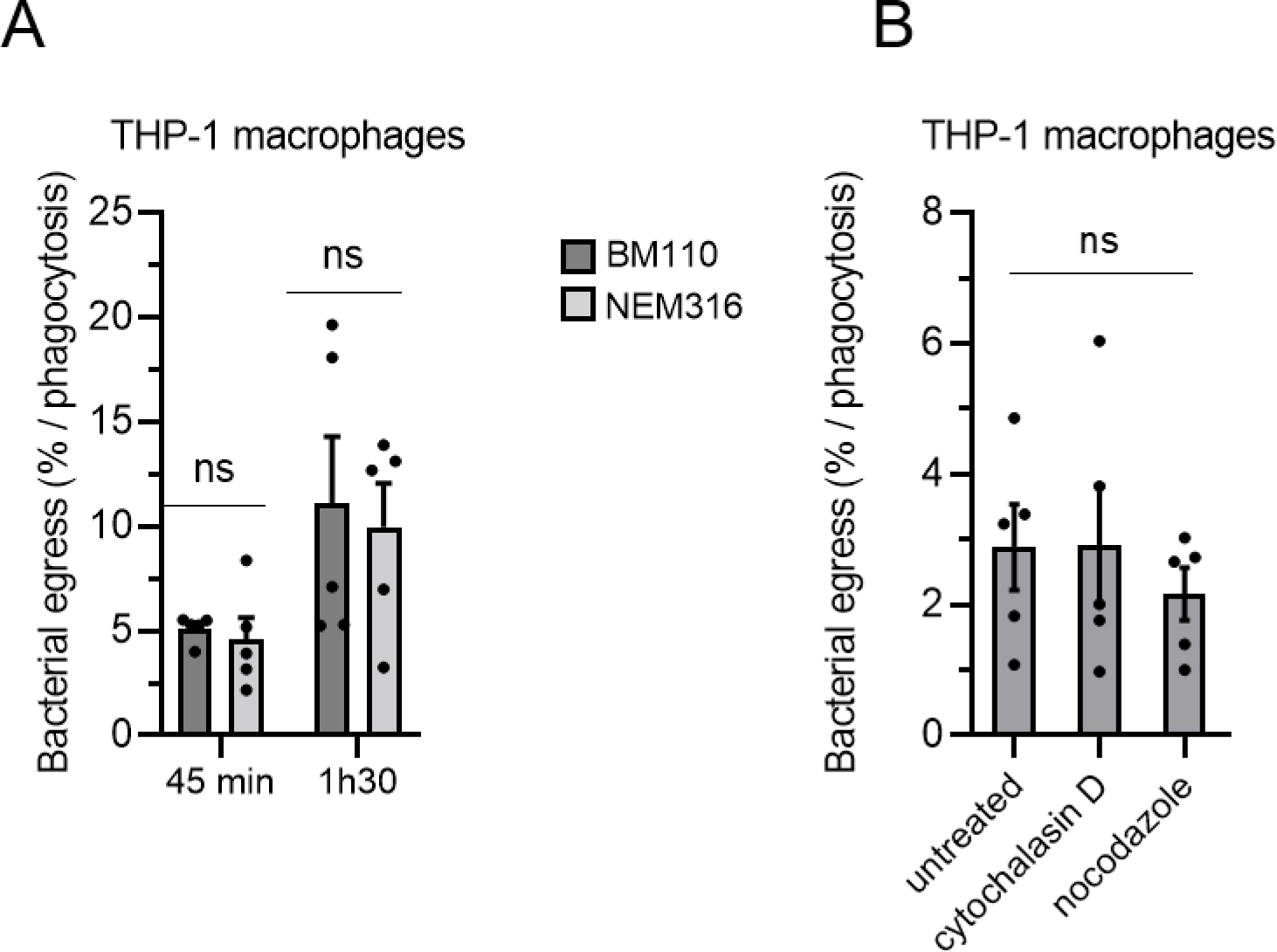
GBS strains can escape macrophages independently of actin and microtubules. (A, B) THP-1 were infected with GBS strains at MOI 10 then treated with antibiotic to kill extracellular bacteria. Bacterial egress was measured by CFU count in cell supernatant devoid of antibiotic after (A) 45 min or 1h30 or (B) 45 min post-antibiotic removal. Results are expressed as the percentage of viable bacteria released in the cell supernatants normalized to the number of phagocytosed bacteria. Bacterial egress was measured for (A) BM110 or NEM316 strains in the absence of treatment or (B) BM110 in the presence of actin (cytochalasin D) or microtubules (nocodazole) depolymerizing agents added after phagocytosis. Statistical analysis: data shown are mean ± SEM of at least four independent experiments. (A) Two-way ANOVA with Dunnett’s multiple comparison (B) one-Way ANOVA with Dunnett’s multiple comparisons tests were performed with ns, non-significant.

In conclusion, both strains of GBS can escape from the intracellular compartment to be released into the extracellular compartment at similar rates. Furthermore, bacterial escape is a passive phenomenon, suggesting that escape does not result from the recycling of the bacteria-containing vacuole to the plasma membrane.

## DISCUSSION

The innate immune response is critical for pathogen clearance, especially in neonates in whom the adaptive immune system is not yet fully mature and the specific antibodies required for adaptive immune response are usually lacking. Phagocytic cells populate all tissues and organs of the body where they play an important role in pathogen clearance.

The ability of GBS lineages to interact specifically with phagocytes, particularly their ability to be phagocytosed, remains controversial. In a study using 35 isolates from various GBS lineages, no significant differences were observed in the phagocytosis of GBS strains stratified by capsular serotype or MLST (30). In contrast, a study performed with a more discrete number of GBS isolates (15 GBS strains including 4 CC17 strains) found that CC17 were phagocytosed by macrophages at significantly higher rates than other GBS isolates (32). Our study, performed with a higher number of invasive GBS clinical isolates (27 CC17 isolates and 40 non-CC17 isolates) demonstrates that GBS CC17 strains are more phagocytosed than non-CC17 strains. These conflicting results may be attributed to the use, in the first study, of heat-inactivation of GBS, that could have denatured bacterial surface proteins involved in phagocytosis (30). In agreement with this hypothesis, we demonstrated that the higher phagocytosis of CC17 strains does not depend on the polysaccharidic capsule, a major GBS virulence factor which protects the bacteria from phagocytosis but, rather on the expression of two surface proteins, Spb1, the backbone protein of the PI-2b pilus, and the HvgA adhesin, that promote a higher level of phagocytosis. In contrast, we did not observe any role for the Srr2 surface protein in GBS CC17 uptake by macrophages. In agreement with this result, we and others have found that blocking the known Srr2-receptors, α5β1 and αvβ3 integrins, by mimetic peptides (RGD and/or RGfV) or by silencing α5 or β3 integrin expression does not affect CC17 phagocytosis (this study and (23)). Together, these data are consistent and indicate that the Srr2 surface protein is not involved in the uptake of GBS by phagocytes.

Importantly, we found that the higher phagocytosis mediated by Spb1 and HvgA was solely the result of an increase of the initial adhesion to phagocytes. Both proteins have previously been described to play a role in adherence to host intestinal epithelial and cerebral endothelial cells (16, 28, 38, 39). Interestingly, in these two cell types, CC17 strains were found to have an approximately 2.5-fold increase in adhesion compared to non-CC17 strains. The increased ability to adhere to both intestinal epithelial and cerebral endothelial cells was found to have physiological functionality, as CC17 strains have a significantly greater ability to colonise the gut and cross the blood-brain barrier (16). Although it is not possible to address *in vivo* whether a 2.3-fold increase in phagocytosis for CC17 strains contributes to the hypervirulence of CC17 strains, our previous data support this hypothesis. After initial adherence to phagocytes, both CC17 and non-CC17 where captured by filopodia and internalized within phagocytes in a process that involved similar signalling molecules. Filopodia formation is known to require the Cdc42 GTPase activity leading to actin polymerization (35). Accordingly, we found that GBS phagocytosis was strongly dependent on actin and Cdc42.

While CC17 had a higher adhesion rate, it shared, with non-CC17 strains, a common signaling pathway leading to phagocytosis. This may reflect the recognition of similar cellular receptors - mediating bacterial adhesion and subsequent internalisation of CC17 and non-CC17 strains - but with different affinities. That both strain types interact with the same molecules but with different affinity has previously been described for GBS interaction with fibrinogen which exhibits a 10-fold higher affinity for CC17 strains than for non-CC17 strains (40). Alternatively, it may reflect an internalisation process in a two-step mechanism involving a *sensus stricto* adhesion receptor followed by engulfment mediated by a phagocytic receptor. In this scenario, CC17 or non-CC17 strains would recognise different adhesion receptors, while the internalisation process would involve a common phagocytic receptor. Two-step phagocytosis mechanisms have previously been described involving cross-talk between TLRs and scavenger receptors (41, 42).

The identity of the cellular receptor(s) involved in the non-opsonized phagocytosis of GBS has been carried out by many teams (22, 23, 43). Despite these efforts, such receptor(s) are still unknown. Our study indicates that FcyRs, complement receptors and CD14 are not involved in the phagocytosis of non-opsonized GBS, as described previously (22, 23, 43). Accordingly, GBS phagocytosis is independent of Syk and Rock kinases that are known to be involved in FCyR and complement receptor signalling, respectively (44). Interestingly, fucoidan and poly(I), two broad scavenger receptor inhibitors reduced the phagocytosis of the BM110 CC17 strain, while having no significant effect on the adhesion and phagocytosis of a non-CC17 strain. Scavenger receptors are a diverse family of proteins that recognize polyanionic ligands and can act as PRR involved in the recognition of various microbial ligands as well as in their phagocytosis (45). The polynucleotide ligand poly(I) acts by efficient competitive inhibition of the binding of other polyanionic ligands to the cationic collagenous binding domain of some scavenger receptors. Similarly, fucoidan easily interacts with positively charged molecules because of its high negative charge at neutral pH (46, 47). These molecules have been successfully used to inhibit adhesion and/or phagocytosis of several pathogens (48, 49). The inhibitory effect of fucoidan and poly(I) on CC17 adhesion and phagocytosis led us to conclude that scavenger receptor(s) might be involved in the adhesion and/or uptake of non-opsonized CC17 bacteria by macrophages. Since scavenger receptors have a wide distribution, being expressed by phagocytes as well as by epithelial and endothelial cells (37), it is tempting to speculate that Spb1 and HvgA promote adhesion to host cells by recognizing the same receptors on phagocytes, epithelial and endothelial cells. Interestingly, it has been recently described that the scavenger receptor SSC4D, expressed by phagocytes and epithelial cells, interacts with BM110 strain (50). SSCD4 is a secreted scavenger receptor making this molecule unlikely to be involved in the initial adherence of CC17 bacteria to the macrophage surface. In addition, silencing SSC4D expression does not affect CC17 phagocytosis (not shown) confirming its absence of involvement in this phenotype. Despite our efforts, the siRNA screen performed to identify scavenger receptor(s) involved in GBS uptake was unsuccessful. Several hypotheses could explain this failure. Because CC17 increased phagocytosis is a phenotype conserved between macrophages and monocytes from cell lines and primary cells, we hypothesized that the receptor(s) involved in CC17 adhesion/phagocytosis is also conserved in these different cell types. However, we cannot exclude the possibility that different scavenger receptors in each cellular model allow CC17 uptake, or that several scavenger receptors have a cooperative effect. Also, the list of scavenger receptors is constantly growing and we might have missed the scavenger receptor(s) involved in GBS CC17 adhesion/phagocytosis. Finally, we cannot exclude a non-specific or an indirect effect of fucoidan and poly(I) on the signaling of others PRRs, as described previously (51).

Numerous pathogens have evolved sophisticated strategies to avoid capture and/or killing by phagocytes. GBS efficiently survive within phagocytes by expressing multiple bacterial factors involved in acid stress response and ROS resistance, including the orange carotenoid pigment hemolysin, the glutathione or the superoxide dismutase (SodA) (26, 52). The capacity of different GBS lineages to survive within macrophages has been controversial (25, 27, 31, 32). The discrepancies between the different studies likely come from either the low number of bacterial strains tested or to the calculation of survival rate (25, 27, 31, 32). Indeed, depending on whether the intracellular survival ability is related to the number of phagocytosed bacteria (intracellular bacteria at T0) or to the initial bacterial inoculum different results will be obtained. As CC17 strains display higher adhesion level leading to higher initial load of intracellular CC17 this might *in fine* lead to a higher recovering number of intracellular bacteria at later time points. By calculating the survival rate in relation to the number of phagocytosed bacteria, we found that CC17 and non-CC17 strains display similar intracellular fate with similar survival capacity. In agreement with this result, we found that Sbp1 and HvgA proteins do not contribute to intracellular survival. We also showed that after internalisation, bacteria can be released into the external compartment. As observed for bacterial survival, we found that bacterial egress of CC17 and non-CC17 strains occurs at a similar rate when normalised to the initially phagocytised bacteria. However, due to enhanced phagocytosis of CC17 strains, this results in a greater number of bacteria being released into the extracellular compartment. Interestingly, we found that bacterial egress occurs even when actin or microtubule networks are disrupted. Since most vacuolar movements involve the cytoskeleton, this suggests that GBS release into the extracellular compartment is a passive phenomenon that may be due to spontaneous macrophage lysis. Further studies are needed to confirm this hypothesis.

The hypervirulent CC17 clone is strongly associated with LOD (60% to 80% of the cases) for which the gastrointestinal tract is likely to be the portal of entry. In the gut, CC17 strains can transcytose *via* M cells and are found associated with CD11c - a marker for macrophages and dendritic cells - positive cells in the Peyer’s patches (8). Interestingly, an increased proportion of CD11c positive cells have been found in the mesenteric lymph node of CC17-infected mice (8). This suggests that CD11c positive cells may act as Trojan horses, to promote the dissemination of CC17 bacteria in the gut. In addition, the higher CC17 intracellular load could also contribute to the establishment of a survival niche and bacterial persistence in tissues.

In conclusion, the specific features associated with the interaction between the CC17 lineage and the phagocytes likely contribute to its particular virulence and to the pathophysiology of infection. Indeed, the increased CC17 bacterial loads in macrophages may confer a selective advantage to promote better persistence and dissemination.

## MATERIALS AND METHODS

### Bacterial strains

Bacterial strains used in this study are listed in Table S1. GBS BM110 (CC17) and NEM316 (CC23) are well characterized capsular serotype III strains isolated from blood cultures of neonates (16). BM110 and NEM316 were used as CC17 and non-CC17 reference strains, respectively. Other GBS clinical strains isolated from invasive infections (*i.e*. from blood culture, cerebrospinal fluid or other normally sterile sites) were obtained from the French National Reference Center for Streptococci (http://www.cnr-strep.fr). Capsular genotyping and MLST was performed for all GBS strains (Table S2). All bacterial strains were cultured in Todd Hewitt (TH) broth or agar (Difco Laboratories, Detroit, MI) at 37°C, except for *L. lactis* which was grown at 30°C. When necessary, antibiotics were used to maintain plasmids at the following concentrations, spectinomycin 100 μg.mL^−1^, erythromycin 10 μg.mL^−1^.

### THP-1 cell line and culture conditions

THP-1 (TIB-202 from ATCC) cell line is a spontaneous immortalized monocyte-like cell line, derived from the peripheral blood of a childhood case of acute monocytic leukemia. THP-1 cells were cultured at 37°C under a 5% CO_2_ atmosphere in RPMI1640 + Glutamax (Gibco), 10% Fetal Calf Serum (FCS, Gibco) and 0.05 mM β-mercapto-ethanol. Cells were passaged twice per week and kept at a cell concentration below 1 × 10^6^ cells /mL. THP-1 cell line was routinely tested for mycoplasma contamination (Mycoalert mycoplasma detection kit, LONZA). To differentiate monocytic THP-1 into adherent macrophages, cells were seeded in 24-well plates at 3 × 10^5^ cells per well in culture media containing 200 nM phorbol 12-myristate 13-acetate (PMA) during 3 days, then media was replaced by fresh RPMI1640 complete media without PMA. Cells were incubated for an additional 5 days without any medium change to allow complete differentiation into macrophages as described previously (53).

### Isolation and culture of primary human Peripheral Blood Mononuclear Cells (PBMCs) and Monocytes-Derived Macrophages (hMDMs)

Blood of adult healthy donors was used (Etablissement Français du Sang, Ile de France, Site Saint-Antoine-le Crozatier) with the appropriate ethics prior approval as stated in the EFS/ INSERM agreement #18EFS030, ensuring that all donors gave a written informed consent and providing anonymized samples. Primary human monocytes were negatively selected using the EasySep Human Monocyte enrichment kit (StemCell®) following the manufacturer’s instructions. For hMDM, primary human PBMCs were isolated by density gradient sedimentation in Histopaque® (Sigma) according to the manufacturer’s recommendations. Monocytes were then selected by adhesion to dishes for 2 h at 37°C in FCS-free medium (RPMI 1640 medium supplemented with 0.1 mg.ml^-1^ gentamicin). Monocytes were differentiated into hMDMs for at least 7 days in complete medium containing 10 ng.mL^-1^ of recombinant human macrophage colony-stimulating factor (rhM-CSF; R&D systems).

### Phagocytosis and survival assay in monocytes and macrophages

Monocytes and macrophages from THP-1 cell line and primary cells were used to study the phagocytosis and the intracellular survival of GBS strains. All experiments were carried out with GBS strains grown to mid-log phase and washed twice with sterile Phosphate buffer saline (PBS) before infection. After addition of bacteria, a quick centrifugation (5 min, 200 × g) was performed to synchronize infection. Phagocytes were infected for 45 min at 37°C and under a 5% CO_2_ atmosphere in RPMI 1640 medium with GBS strains, using a multiplicity of infection (MOI) of 10 bacteria per cell except when otherwise specified. After infection, phagocytes were washed 3 times with PBS. Remaining extracellular bacteria were killed using RPMI 1640 medium supplemented with Penicillin G / Streptomycin (100 units.mL^-1^ / 0.1 mg.mL^-1^) and Gentamicin (0.2 mg.mL^-1^) at 37°C and 5% CO_2_. After antibiotic treatment, cells were washed twice and either lysed directly by osmotic choc using sterile ice-cold H_2_O to evaluate phagocytosis (T0) or further incubated in RPMI1640 medium to follow intracellular survival before being lysed (2h30 or 24h). Serial dilutions of the lysates were plated onto TH agar for colony forming units (CFU) counting. The percentage of phagocytosis was calculated as follows: (CFU phagocytosis / CFU in inoculum) × 100. The percentage of survival was calculated as follows: (CFU survival / CFU phagocytosis) × 100. When specified, results were expressed as normalized to the control strain (BM110). Assays were performed in triplicate and were repeated at least three times.

### Adhesion assay onto monocytes and macrophages

For adhesion assays, infection was performed as described for phagocytosis but at 4°C and with ice-cold RPMI1640 medium to avoid engulfment of bacteria and phagocytosis. In some experiments, adhesion was also tested at 37°C using cells that had been pre-treated for 1 hour with cytochalasin D. After 3 washes with ice-cold PBS to remove non-adherent bacteria, phagocytes were directly lysed by osmotic shock with ice-cold H_2_O. Serial dilutions of lysates were then plated onto TH agar for CFU counting. The percentage of adhesion was determined by calculating the ratio between the CFUs of adherent bacteria and the CFUs from the inoculum × 100. When specified, results were expressed as normalized to the control strain. Assays were performed in triplicate and were repeated at least three times.

### Phagocytosis assay in the presence of inhibitors/reagents

Selected compounds were tested for their ability to block GBS phagocytosis. All inhibitors and reagents used in this study are listed in Table S3. THP-1 macrophages were pretreated with inhibitors/reagents for 1h prior infection with BM110 or NEM316 strains. Inhibitors/reagents were left during the time of infection at the same concentration. Treatments did not affect bacterial or cell viability as assessed by Lactate dehydrogenase release in the culture supernatant (LDH cytotoxicity kit^®^, Promega, according to the manufacturer’s instructions; data not shown). The percentage of phagocytosis was calculated as follows: (CFU phagocytosis / CFU in inoculum) × 100 and results were normalized to the untreated condition.

### Evaluation of bacterial egress from macrophages

To assess bacterial egress from macrophages, cells were infected as described previously, and then treated with antibiotics to kill extracellular bacteria for 45 min. Cells culture medium was next replaced by medium without any antibiotic. Bacterial egress was quantified in the supernatant of infected macrophages after 45 min or after 1h30 following the replacement of the cell culture medium. The percent of bacterial egress was calculated as follows: (CFU released bacteria / CFU phagocytosed) × 100.

### RNA extraction, cDNA synthesis and RT-PCR

RNAs were extracted from non-infected THP-1 macrophages, monocytes and hMDM using the NucleoSpin® RNA kit (Macherey-Nagel®) according to the manufacturer’s instructions. RNA concentration was determined using a NanoDrop 2000 spectrophotometer. Then, 1 μg RNA was reverse-transcribed into cDNA using the SuperScript® IV Reverse Transcription Kit (Invitrogen), following the manufacturer’s instructions. PCR were carried out with Green Taq DNA polymerase (Promega) on cDNA using primers described in Table S4. Primers were designed in order to amplify all variants of the targeted receptor(s). Amplification products were resolved in 1% (w/v) agarose gel.

### siRNA transfection

To silence the expression of putative phagocytic receptors, pools of three or four siRNA duplexes (Dharmacon) were used. The silencer select-negative control-1 siRNA (Ambion) was used as negative control. THP-1 were seeded in twenty-four-well plates for 24 h in the presence of 200 nM of PMA then transfected with 50 nM siRNA using Lipofectamine RNAIMAX (Invitrogen) according to the manufacturer’s instructions and incubated for a further 3 days. Efficiency of knockdown was assessed by Western Blot analysis on 2 independent targets.

### Immunofluorescent staining and image analysis

To follow intracellular fate of GBS strains, THP-1 macrophages were infected for 45 minutes as described for phagocytosis then washed extensively and fixed at different time points post-phagocytosis (0, 2h30 or 5h) on glass coverslips with 4% paraformaldehyde (PFA) for 10 min then washed with PBS. Following fixation, samples were permeabilized and saturated with PBS containing 0.2% saponin and 10% serum for 1h at room temperature, then incubated for at least 1h at room temperature with the primary anti-LAMP1 antibody (DSHB clone H4A3). After 3 washes with PBS, samples were incubated for 1h at room temperature with conjugated-secondary antibodies (Jackson ImmunoResearch Laboratories).

To enable detection of acidic compartments (phagolysosome), THP-1 macrophages were incubated 30 min before the end of infection with Lysotracker red DND-99 (Invitrogen) at a final concentration of 75 nM, fixed with 4% PFA for 10 min, then washed with PBS.

After 3 washes in PBS, coverslips were mounted in DAKO fluorescent mounting media (DAKO). Image acquisitions were realized using confocal laser scanning microscopy (Leica DMI6000) coupled to a spinning disk confocal head (YokogawaCSU-X1M1), controlled by the Metamorph 7.7.5 software (Cochin Institute microscopy facility, IMAG’IC). Images were processed using the ImageJ software. At least 300 cells were analyzed for each sample in four independent experiments.

### Scanning Electron Microscopy

All incubations were performed at room temperature except when otherwise indicated. Macrophages grown on glass coverslips were infected as described previously and then fixed in 0.1M sodium cacodylate buffer, pH 7.4 containing 2.5% glutaraldehyde and 1% PFA for 60 min. After washing in cacodylate buffer (2×10 min), cells were fixed in 1% OsO_4_ diluted in cacodylate buffer for 45 min at 4°C. After washing in cacodylate buffer (2×10 min), samples were dehydrated in an ascending series of ethanol (30%, 50%, 70%, 95%, 100%, 100%, 100% - 10 min each), followed by hexamethyldisilazane /ethanol (1/1) for 10 min and hexamethyldisilazane for 10 min. After overnight air drying, each coverslip was placed on a double-sided sticky tape on the top of an aluminum stub and sputter coated with Au/Pd. Images were acquired using a Jeol LV6510 (Jeol, Croissy-sur-Seine, France).

### Statistical analysis

All assays were performed in at least in three independent experiments, each experiment was carried out in triplicate. Data represent mean ± SEM or median and statistical analyses were performed using GraphPad Prism 10.0 (GraphPad Software, San Diego, California). Significance levels were set at \**p* ≤ 0.05; \*\**p* ≤ 0.01; \*\*\**p* ≤ 0.001; **** *p* ≤ 0.0001.

## AUTHOR CONTRIBUTIONS

JG designed the research. ASB, AP, JCF, CH and JG performed the experiments. ASB, AP, JG analyzed the data. BS and VM performed electron microscopy experiments. AF, AT, CP provided intellectual input and guidance. JG wrote the manuscript and ASB, AF, AT contributed to finalizing the manuscript. All authors discussed results and reviewed the final version of the manuscript.

## ACKNOWLEDGMENTS

We are grateful to Glen Ulett for the gift of the GFP plasmid. We thank Cristina Candeias for technical assistance, Shaynoor Dramsi for BM110Δ*spb1* and NEM316Δ*cps* mutant strains and the Imag’IC core facility of the Cochin Institute. The H4A3 antibody developed by August, J.T. / Hildreth, J.E.K. was obtained from the Developmental Studies Hybridoma Bank, created by the NICHD of the NIH and maintained at The University of Iowa, Department of Biology, Iowa City, IA 52242.

## Abbreviations

CC17: clonal complex 17
GBS: group B *Streptococcus*)

