## supplementary file for "Specific interaction between Group B *Streptococcus* CC17 hypervirulent clone and phagocytes"

**TABLE S1**: Bacterial strains used in this study.

| Strains | Relevant characteristics | Reference |
|---|---|---|
| BM110 | serotype III, CC17, bacteremia | (1) |
| BM110-GFP | BM110 strain carrying pGU2664 plasmid for GFP expression | (1) |
| Δ*srr2* | BM110 strain deleted in *srr2* gene | (2) |
| Δ*hvgA* | BM110 strain deleted in *hvgA* gene | (3) |
| Δ*spb1* | BM110 strain deleted in *spb1* gene | (4) |
| BM110 Δ*cps* | BM110 strain deleted in *cps*D/E genes (capsular mutant) | (5) |
| NEM316 | serotype III, CC23, EOD bacteremia | (3) |
| NEM316-GFP | NEM316 strain carrying pGU2664 plasmid for GFP expression | This study |
| NEM316 Δ*cps* | NEM316 strain deleted in *cps*D/E gene (capsule mutant) | (5) |
| *L. lactis* + vector | MG1363 strain of *Lactococcus lactis* subsp. *cremoris* carrying empty pOri232 plasmid | (3) |
| *L. lactis* + HvgA | MG1363 strain of *Lactococcus lactis* subsp. *cremoris* carrying pOri232 plasmid for HvgA expression | (3) |

**TABLE S2:** GBS clinical isolates used in this study.

| Strains | Capsular serotype | Complex clonal (CC) | Type of infection |
|---|---|---|---|
| CNR CCH1569 | CPS III | CC17 | LOD Meningitis |
| CNR CCH1570 | CPS III | CC17 | LOD Meningitis |
| CNR CCH1571 | CPS III | CC17 | LOD Meningitis |
| CNR CCH1573 | CPS III | CC17 | LOD Bacteremia |
| CNR CCH1575 | CPS III | CC17 | LOD Meningitis |
| CNR CCH1577 | CPS III | CC17 | EOD Meningitis |
| CNR CCH1578 | CPS III | CC17 | LOD Meningitis |
| CNR CCH1581 | CPS III | CC17 | EOD Meningitis |
| CNR CCH1584 | CPS III | CC17 | LOD Meningitis |
| CNR CCH1586 | CPS III | CC17 | LOD Meningitis |
| CNR CCH1588 | CPS III | CC17 | LOD Meningitis |
| CNR CCH1589 | CPS III | CC17 | LOD Meningitis |
| CNR CCH1591 | CPS III | CC17 | LOD Meningitis |
| CNR CCH1594 | CPS III | CC17 | LOD Meningitis |
| CNR CCH1596 | CPS III | CC17 | LOD Meningitis |
| CNR CCH1597 | CPS III | CC17 | LOD Meningitis |
| CNR CCH1602 | CPS III | CC17 | EOD Meningitis |
| CNR CCH0620 | CPS IV | CC17 | Adult Bacteremia |
| CNR CCH0755 | CPS IV | CC17 | Adult Bacteremia |
| CNR CCH0778 | CPS IV | CC17 | Adult Bacteremia |
| CNR CCH0950 | CPS IV | CC17 | Adult Bacteremia |
| CNR CCH1030 | CPS IV | CC17 | Adult Meningitis |
| CNR CCH1174 | CPS IV | CC17 | Adult Bacteremia |
| CNR CCH1239 | CPS IV | CC17 | Adult Bacteremia |
| CNR CCH1360 | CPS IV | CC17 | Adult Meningitis |
| CNR CCH1472 | CPS IV | CC17 | EOD pneumonia |
| CNR CCH1506 | CPS IV | CC17 | LOD Bacteremia |
| CNR CCH0050 | CPS III | non-CC17 | EOD Bacteremia |
| CNR CCH0107 | CPS III | non-CC17 | LOD Meningitis |
| CNR CCH0112 | CPS III | non-CC17 | EOD Bacteremia |
| CNR CCH0150 | CPS III | non-CC17 | LOD Bacteremia |
| CNR CCH0205 | CPS III | non-CC17 | LOD Bacteremia |
| CNR CCH0311 | CPS III | non-CC17 | EOD Bacteremia |
| CNR CCH0362 | CPS III | non-CC17 | LOD Bacteremia |
| CNR CCH0382 | CPS III | non-CC17 | Adult osteoarticular infection |
| CNR CCH0384 | CPS III | non-CC17 | LOD Bacteremia |
| CNR CCH0513 | CPS III | non-CC17 | LOD Meningitis |
| CNR CCH0700 | CPS III | non-CC17 | LOD Meningitis |
| CNR CCH1207 | CPS III | non-CC17 | LOD Meningitis |
| CNR CCH1261 | CPS III | non-CC17 | LOD Bacteremia |
| CNR CCH1280 | CPS III | non-CC17 | LOD Meningitis |
| CNR CCH1319 | CPS III | non-CC17 | LOD Bacteremia |
| CNR CCH1393 | CPS III | non-CC17 | LOD Bacteremia |
| CNR CCH1396 | CPS III | non-CC17 | LOD Bacteremia |

| | | | |
|---|---|---|---|
| CNR CCH1490 | CPS III | non-CC17 | Intra-uterine infection (pregnancy-associated) |
| CNR CCH1809 | CPS III | non-CC17 | LOD Bacteremia |
| CNR CCH1881 | CPS III | non-CC17 | LOD Meningitis |
| CNR CCH1585 | CPS IV | non-CC17 | EOD Bacteremia |
| CNR CCH0966 | CPS IV | non-CC17 | EOD Meningitis |
| CNR CCH1134 | CPS IV | non-CC17 | Adult Bacteremia |
| CNR CCH1157 | CPS IV | non-CC17 | LOD pneumonia |
| CNR CCH1165 | CPS IV | non-CC17 | EOD Bacteremia |
| CNR CCH1274 | CPS IV | non-CC17 | Adult Bacteremia |
| CNR CCH1363 | CPS IV | non-CC17 | LOD Bacteremia |
| CNR CCH1369 | CPS IV | non-CC17 | Adult Bacteremia |
| CNR CCH1403 | CPS IV | non-CC17 | Adult Bacteremia |
| CNR CCH1686 | CPS IV | non-CC17 | LOD Bacteremia |
| CNR CCH1565 | CPS II | non-CC17 | EOD Bacteremia |
| CNR CCH1566 | CPS Ia | non-CC17 | EOD Meningitis |
| CNR CCH1567 | CPS Ia | non-CC17 | EOD Bacteremia |
| CNR CCH1568 | CPS Ia | non-CC17 | EOD Bacteremia |
| CNR CCH1572 | CPS Ia | non-CC17 | EOD Bacteremia |
| CNR CCH1576 | CPS Ia | non-CC17 | EOD Bacteremia |
| CNR CCH1579 | CPS V | non-CC17 | EOD Bacteremia |
| CNR CCH1580 | CPS V | non-CC17 | LOD Bacteremia |
| CNR CCH1582 | CPS Ib | non-CC17 | LOD Meningitis |
| CNR CCH1583 | CPS Ia | non-CC17 | LOD Bacteremia |

**TABLE S3:** Inhibitors used in this study.

| Chemical and reagents | Target | Concentration |
|---|---|---|
| Cytochalasin D | Actin filaments | 2 µM |
| Nocodazole | Microtubules | 1 µM |
| LY29002 | Pi 3-kinase | 40 µM |
| PP2 | Src kinase | 10 µM |
| Staurosporin | Protein kinase C | 500 nM |
| NSC23766 | Rac | 100 µM |
| ZCL278 | CDC42 | 50 µM |
| Bay61-3606 | Syk | 1 µM |
| Y27632 | Rock | 10 µM |
| Human FC Block™ | FCy receptors | 2 µg/ml |
| Anti CR3 antibody (vim12) | Complement Receptor 3 | 20 µg/ml |
| IgG1 mouse antibody | Isotype control | 20 µg/ml |
| Mannan | Lectin receptors | 200 µg/ml |
| Mannose | Lectin receptors | 10 mg/ml |
| Laminarin | Lectin receptors | 200 µg/ml |
| RGD peptide | RGD dependent integrins | 50 µM |
| RGDfV peptide | RGDfV dependent integrins | 50 µM |
| Fucoidan | Scavenger receptors | 500 µg/ml |
| Poly (I) | Scavenger receptors | 200 µg/ml |
| Poly (C) | Inactive analog of Poly(I) | 200 µg/ml |

**TABLE S4:** Primers used in this study.

| Target | Primers | Sequence (5'-3') | Expected size (bp) |
|---|---|---|---|
| Actin | F-1462<br>R-1580 | TTCCAAATATGAGATGCGTTGTTA<br>ATGCTATCACCTCCCCTGTG | 118 |
| VE-cadherin | F-453<br>R-571 | GGTCGATGCAGAGACAGGAG<br>GAGTCTCCAGGTTTTCGCCA | 118 |
| SR-A1/MSR1 | F-595<br>R-707 | GCCAACCTCATGGACACAGA<br>CCATGTCCCTGGACTGAGGA | 112 |
| SR-A6/MARCO | F-1228<br>R-1361 | TGTGGAGCTGCACCAAGAAT<br>CCACATATGAGCCCGAGGAC | 133 |
| SR-A3/SCARA3 | F-866<br>R-992 | CGAGGAGACCCTGACCCTCCAG<br>CCCAGGGTGGCCTGGATGTTC | 126 |
| SR-A4/SCARA4 | F-618<br>R-796 | ACGCTGGAGAAGTTACAGGC<br>TTCTGCAGATTGCCCTGGAG | 178 |
| SR-A5/SCARA5 | F-2386<br>R-2579 | CCATGCACCAGGCCTCAATA<br>CACTTGACGTTGCCTCTTGC | 193 |
| SR-B2/CD36 | F-1559<br>R-1682 | CAATTTGCAAAACGGCTGCAG<br>CTTCTCATCACCAATGGTCCC | 123 |
| SR-B1/SCARB1 | F-1520<br>R-1652 | AGGGGGAGACTCTTCACACA<br>GGCTCCGGATTTGGCAGATG | 132 |
| SR-D1/CD68 | F-1093<br>R-1261 | GATCCTTCTTGGCCTCCTCG<br>CTTTGAGCCAGTTGCGTGTC | 168 |
| SR-E1/LOX1 | F-740<br>R-920 | CGAGGAGCTGTTTATGCGGA<br>TGGCACCCAAGTGACAAAGA | 180 |
| SR-E2/Dectin1 | F-830<br>R-943 | ACCCATCTCCAAATTGTGTA<br>CCACCCTTCCTCTTACATTG | 113 |
| SR-F1/SCARF1 | F-2964<br>R-3100 | CTCCCTCTGTCCCCCAGGCT<br>GCCAAGCGTGGTGGAGGGCACC | 136 |
| SR-G1/CXCL16 | F-1041<br>R-1162 | CCACCCTCCCAGTAGGATCA<br>CTGCTTCTGGTTCTCCCCAG | 121 |
| SR-H1/FEEL1 | F-774<br>R-937 | CAACCCCTGCTGGCCATCAC<br>GTAGAGTTGCTGGGGCAGCC | 163 |
| SR-H2/FEEL2 | F-4194<br>R-4375 | GCTGTGCCGGCTTCTTTGGC<br>GGTCACAGTGGATGCCGTAC | 181 |
| SR-I1/CD163 | F-310<br>R-455 | GCGGGAGAGTGGAAGTGAAA<br>ACCTGCACTGGAATTAGCCC | 145 |
| DC-SIGN/CD209<br>DC-SIGN/CD209 | F-2342<br>R-2484 | GAATTGTGCCTCTCCTGGCT<br>GTGGGCCACCACGATGAATA | 142 |
| SR-E4/ASGPR | F-155<br>R-289 | GGAATCAGACCCTGAGACCC<br>TGCAGCTGGGAGTCTTTTCTG | 134 |
| SR-F2/MEGF10 | F-29<br>R-177 | AGCAAGTACTTTCCCGGTGC<br>CAATCAACGCGGTTAGCGTC | 148 |
| SR-J1/RAGE | F-1431<br>R-1571 | GGCCCACAGACAGATCCCAT<br>GGGGGCTCTGGTTGTAGAAG | 140 |
| SR-L1/LRP1 | F-3616 | GACTGTGGGGACAACAGTGAC | 154 |

| | | | |
|---|---|---|---|
| | R-3770 | TTGGCGTGTGTCTCATCACT | |
| SR-L2/LRP2 | F-15028 | AGCCTCAACTGGGTTTTTGT | 129 |
| | R-15157 | GTACACATTTAGCCACAGGGC | |
| SSC4D/SRCRB4D | F-2046 | GGACAATGTCAAGTGCCGTGGG | 182 |
| | R-2228 | GTAGTCATCACAAGAGGAGGGC | |
| SSC5D | F-1990 | CTGGAGAAAACAACCACGAAG | 170 |
| | R-2160 | GTCAGTGCTGCAGTGGTCTGTG | |
| CD14 | F-352 | CTGACACGGTCAAGGCTCTC | 114 |
| | R-466 | AGTTCCTTGAGGCGGGAGTA | |
| LY75 | F-5058 | ACTGATGAGAAACCCGCCAG | 127 |
| | R-5185 | ACTGATGAGAAACCCGCCAG | |
| CD207/Langerin | F-963 | ATGCCCCATGTGACAAAACG | 103 |
| | R-1066 | GCGTTGGAGCTCAAAGAGTG | |

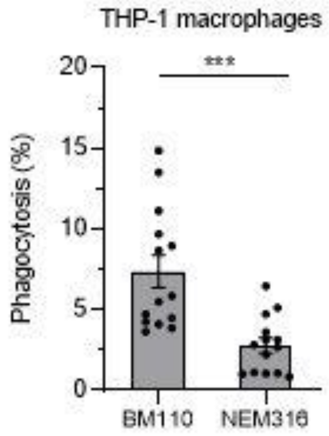


**Figure S1: BM110 are more phagocytosed than NEM316 in THP-1 macrophages.** Phagocytosis level of BM110 and NEM316 strains was assessed by CFU count after infection at MOI 10 followed by antibiotic treatment to kill extracellular bacteria. Results are expressed as the percentage of phagocytosed bacteria normalized to the initial inoculum.

Statistical analysis: data shown are mean ± SEM of at least four independent experiments. t test was performed with ***, $p < 0.001$.

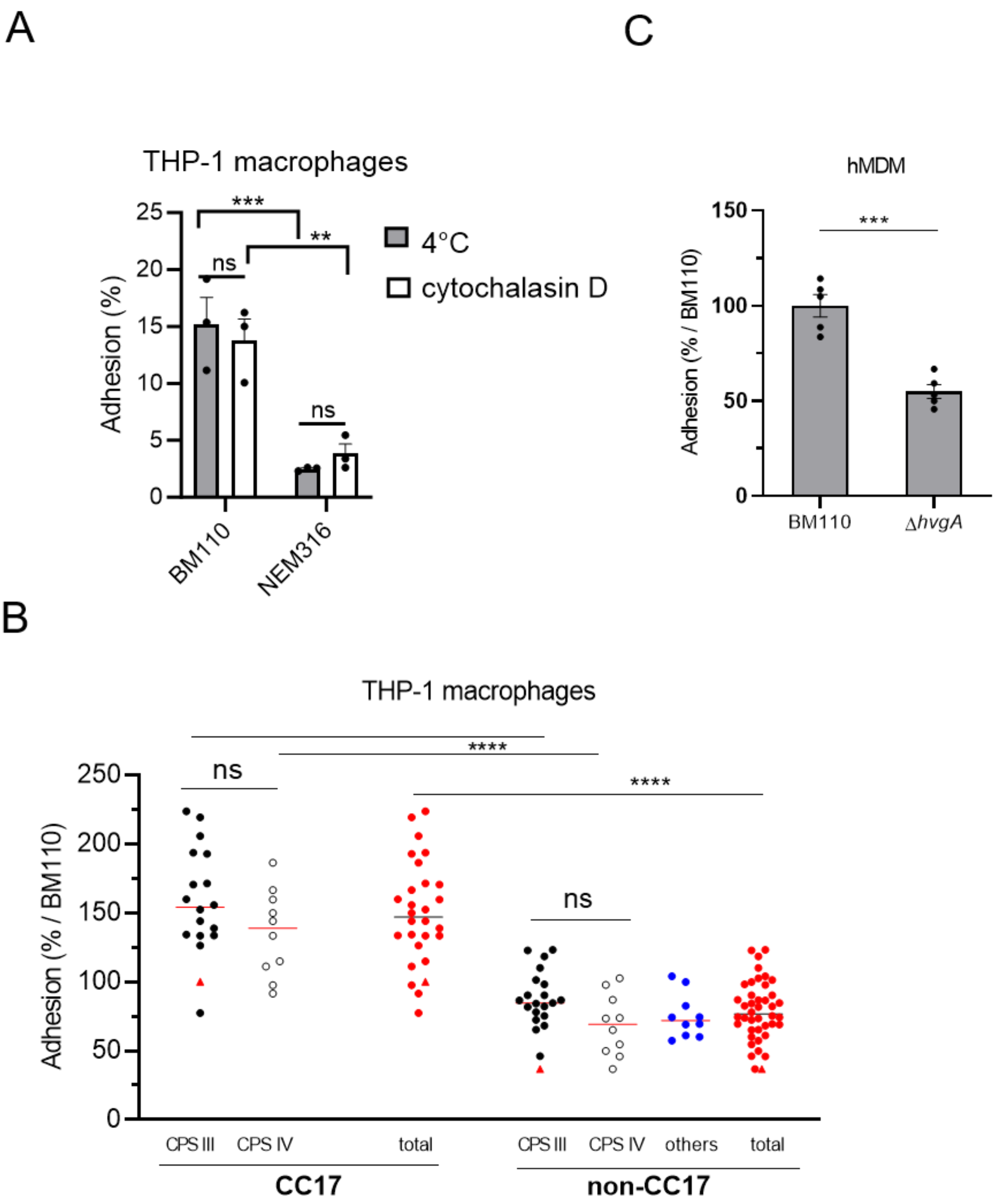


**Figure S2: GBS adhesion to phagocytes.** (A) Adhesion level of GBS strains was assessed by CFU count after infection at MOI 10 at (A) 37 °C in the presence of cytochalasin D or at (A, B, C) 4°C to avoid bacterial engulfment. Adhesion was assessed in (A, B) THP-1 macrophages or (C) hMDM. (A) Adhesion level of BM110 and NEM316 strains are expressed as the percentage of adherent bacteria normalized to the initial inoculum. (B) Adhesion level of CC17 and non-CC17 GBS clinical isolates from invasive infections, each point representing a clinical strain. Triangles correspond to BM110 strain (CC17) and NEM316 (non-CC17). Results are shown according to capsular type (CPS III, CPS IV, or other CPS) and are expressed as the percentage of adhesion relative to BM110 strain adhesion, with horizontal lines indicating median value. (C) Adhesion level of BM110 and its derivative mutant strain (Δ*hvgA*). Results are expressed as the percentage of adherent bacteria normalized to BM110 strain adhesion.

Statistical analysis: data shown are mean ± SEM of at least three independent experiments. (A, B) Two-way ANOVA or (C) t test, were performed with ns, non-significant; ***, $p < 0.001$; ****, $p < 0.0001$.

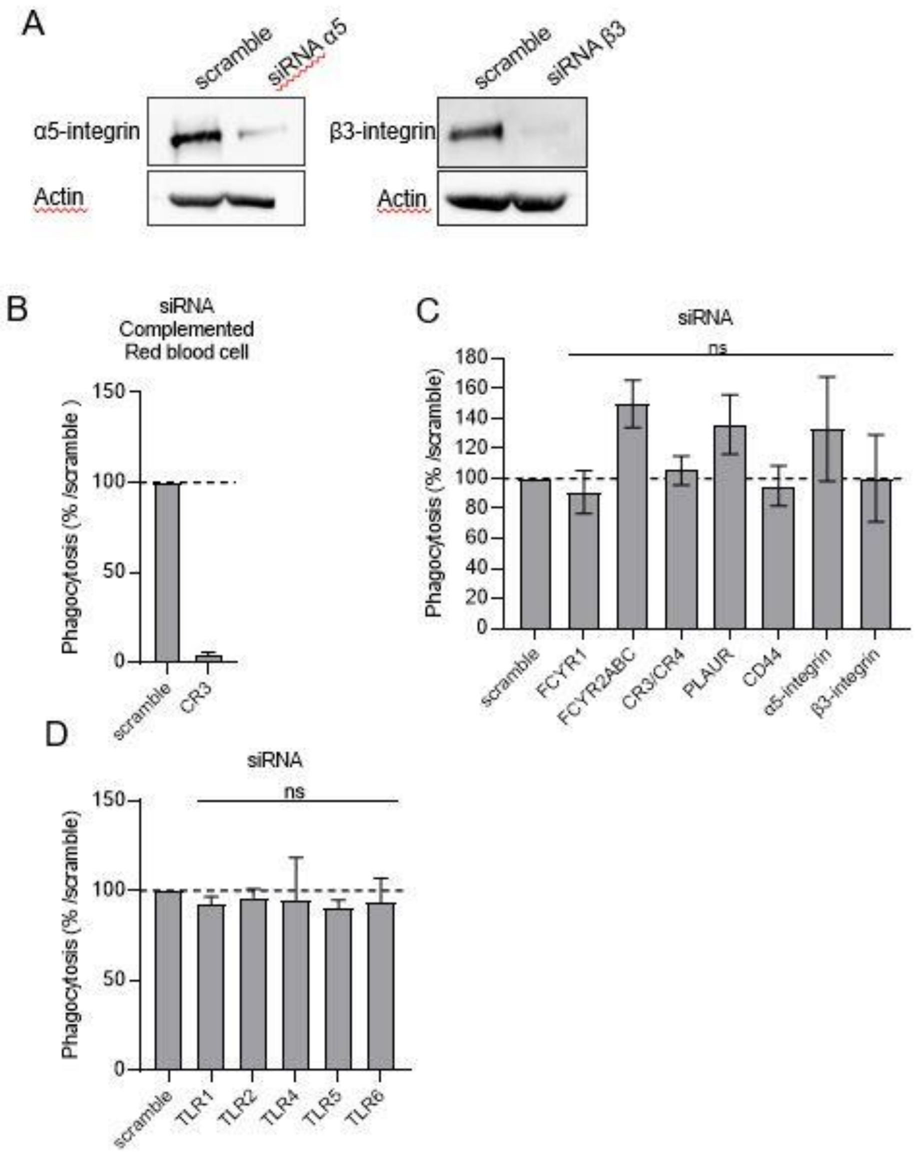


**Figure S3: siRNA screen in THP-1 macrophages.** (A) Silencing efficiency obtained by siRNA treatment of THP-1 macrophages was evaluated by western blot analysis on two independent targets (integrin α5 and β3). (B) Phagocytosis of complement-opsonized red blood cells was evaluated my fluorescence microscopy in THP-1 treated with scramble siRNA or siRNA targeting complement receptor 3(CR3, *itgaM*). Results are expressed as the percentage of phagocytosed red blood cells normalized to scramble control condition. (C, D) Phagocytosis of BM110 strain was evaluated in THP-1 macrophages treated with (C) siRNA targeting FCγR1; FCγR2A,B,C; Complement Receptor3 and 4 (CR, *itgAM*), PLAUR, SR-K1 (CD44) and integrins α5 and β3 or (D) TLRs receptors. Results are expressed as the percentage of phagocytosis normalised to the scramble control conditions.

Statistical analysis: data shown are mean ± SEM of at least three independent experiments. (C, D) Kruskall Wallis test with Dunn’s multiple comparisons tests were performed with ns, non-significant.

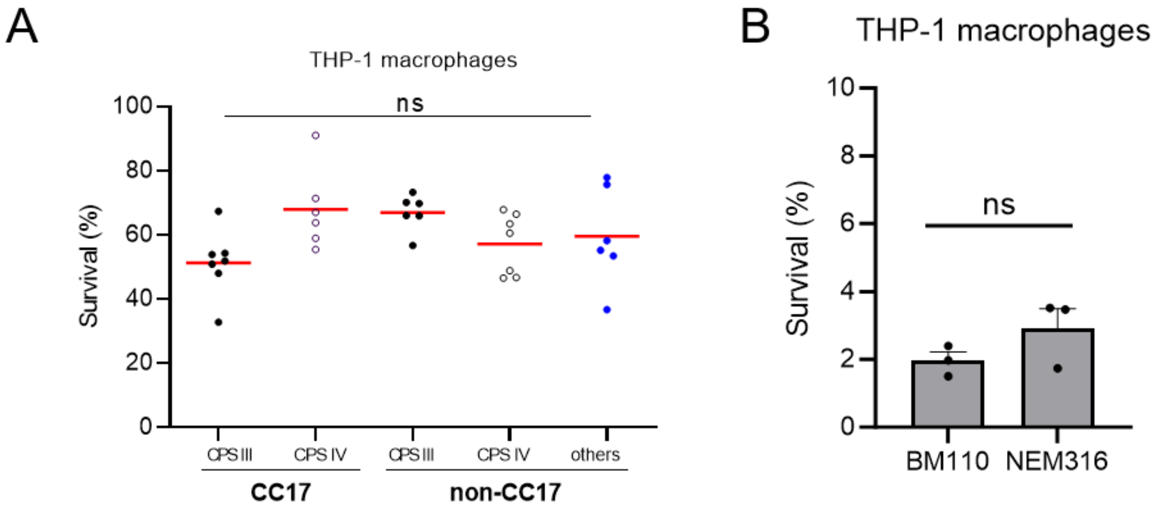


**Figure S4: Survival of GBS strains.** Survival level of GBS strains in THP-1 macrophages was assessed by CFU count. Results are expressed as the percentage of viable intracellular bacteria normalised to the number of phagocytosed bacteria. (A) Survival level of CC17 and non-CC17 GBS clinical isolates from invasive infections was determined 2.5 h after phagocytosis. Each point represents one clinical strain. Results are shown according to capsular type (CPS III, CPS IV, or other CPS). (B) Survival level of BM110 and NEM316 strains was determined 24 h post-phagocytosis.

Statistical analysis: data shown are mean ± SEM of at least three independent experiments. Two-way ANOVA was performed with ns, non-significant.
